# Genome-wide association studies across environmental and genetic contexts reveal complex genetic architecture of symbiotic extended phenotypes

**DOI:** 10.1101/2021.08.03.454976

**Authors:** Rebecca T. Batstone, Hanna Lindgren, Cassandra M. Allsup, Laura A. Goralka, Alex B. Riley, Michael A. Grillo, Amy Marshall-Colon, Katy D. Heath

## Abstract

A goal of modern biology is to develop the genotype-phenotype (G→P) map, a predictive understanding of how genomic information generates trait variation that forms the basis of both natural and managed communities. As microbiome research advances, however, it has become clear that many of these traits are symbiotic extended phenotypes, being governed by genetic variation encoded not only by the host’s own genome, but also by the genomes of myriad cryptic symbionts. Building a reliable G→P map therefore requires accounting for the multitude of interacting genes and even genomes involved in symbiosis. Here we use naturally-occurring genetic variation in 191 strains of the model microbial symbiont *Sinorhizobium meliloti* paired with two genotypes of the host *Medicago truncatula* in four genome-wide association studies to study the genomic architecture of a key symbiotic extended phenotype – partner quality, or the fitness benefit conferred to a host by a particular symbiont genotype, within and across environmental contexts and host genotypes. We define three novel categories of loci in rhizobium genomes that must be accounted for if we want to build a reliable G→P map of partner quality; namely, 1) loci whose identities depend on the environment, 2) those that depend on the host genotype with which rhizobia interact, and 3) universal loci that are likely important in all or most environments.

**IMPORTANCE:** Given the rapid rise of research on how microbiomes can be harnessed to improve host health, understanding the contribution of microbial genetic variation to host phenotypic variation is pressing, and will better enable us to predict the evolution of (and select more precisely for) symbiotic extended phenotypes that impact host health. We uncover extensive context-dependency in both the identity and functions of symbiont loci that control host growth, which makes predicting the genes and pathways important for determining symbiotic outcomes under different conditions more challenging. Despite this context-dependency, we also resolve a core set of universal loci that are likely important in all or most environments, and thus, serve as excellent targets both for genetic engineering and future coevolutionary studies of symbiosis.

## INTRODUCTION

We live in a symbiotic world. It is increasingly recognized that many important traits, including human metabolism, insect diet and defense, and plant nutrient foraging, are actually *symbiotic extended phenotypes* governed (at least in part) by cryptic variation in their microbial symbionts (e.g., 1, 2, 3, 4, 5, 6, 7; reviewed by 8, 9, 10, 11, 12). Incorporating the complex genetics arising from these interactions, however, has lagged behind understanding their ecological outcomes. Characterizing the genetic basis of these cross-domain relationships represents a symbiotic extension of the genotype-phenotype (G→P) map (13, 14, 15), wherein genetic variation present in symbionts (G) presents as phenotypic variation (P) expressed by the host. In symbiosis, as in all organisms, the G→P map is a crucial step towards a predictive understanding of evolution (e.g., a trait’s genomic architecture influences the rates and trajectories of its response to selection), and for bioengineering, where we must know not only which genes to target and exactly how to edit them, but also anticipate how consistent the knock-on effects will be across diverse environments and/or genetic backgrounds (16, 17).

A key lesson from studies of the G→P map within single organisms is that phenotypic variation fundamentally arises in non-additive ways as genomic variation is filtered through “layer(s) of context-dependence” (14) including genotype-by-environment (G × E) interactions, epistasis, and pleiotropy (15, 18, 19, 20, 21, 22, 23). The role of context-dependency in the G→P is particularly relevant for symbiotic extended phenotypes, given the potential for higher-order interactive genetic effects. For example, interacting genomes can generate genotype-by-genotype interactions between host and symbiont (intergenomic epistasis, or G × G: 24, 25, 26, 27, 28, 29) and even G × G × E whereby trait values depend not only on the interaction of alleles in both partners, but on the environmental context (30, 31, 32). Such complex context-dependent effects are critical for evolution; for example, G × G × E for fitness is the statistical expression of the coevolutionary selection mosaic (27, 33, 34). Additionally, genes that consistently affect symbiotic extended phenotypes arising in plant-microbe interactions, independent of these layers of context dependency, are important targets for breeding and bioengineering (9, 16, 35, 36, 37). Fine-scale mapping of the loci contributing to symbiotic extended phenotypes across partner genotype and environmental contexts using multiple genome-wide association studies (GWAS) can therefore generate a more holistic picture of both types of genetic effects (i.e., the genes that universally contribute and those that are context-dependent), and thus, provides insights both for natural (co)evolutionary processes and sustainable agriculture.

The symbiosis between nitrogen-fixing rhizobia and leguminous plants has been a key model for addressing both additive and interactive genetic effects in symbiosis (i.e., G × G, G × E: 38, 39, 40, 41, 42), with resequencing projects, germplasm collections, and many genetic and genomic tools, particularly in *Medicago-Sinorhizobium* interactions (43, 44, 45, 46, 47). These symbioses are also major drivers of the terrestrial nitrogen (N) cycle, contributing up to 21 million tonnes of fixed N each year (48, 49). In their specialized root nodules, legumes trade photosynthetically-derived fixed carbon in exchange for N fixed by the rhizobia, making legumes keystone members of natural plant communities and sustainable agriculture (reviewed by 50, 51). Rhizobial genomes themselves have been studied as models of bacterial genome evolution, reflecting the dynamic tension between the core set of genes, often on the main chromosome, and the more flexible genes, often found on mobile plasmids, islands, and other elements that contain the canonical symbiosis genes (i.e., nod, fix, nif; 52, 53, 54, 55, 56). Symbiosis plasmids often show abundant recombination (45, 57, 58), allowing genome-wide association studies (GWAS) to detect associations between individual genomic variants and traits of interest (45, 56, 59, 60, 61). This symbiosis is thus well poised for understanding the genetic basis of important symbiotic extended phenotypes.

Decades of functional genetic studies have resolved several symbiosis genes, mostly those that disrupt symbiosis when knocked out (reviewed by 62, 63), enabling increasingly well-resolved models for how interactions establish and how trade is coordinated (64). Nevertheless, GWAS in legume-rhizobium symbiosis, which leverage standing genetic variation, indicate that a much broader set of genes and genetic pathways contribute to the quantitative variation in traits that we know to be important in natural and managed systems (45, 46, 59, 65, 66), especially **partner quality** - or the fitness benefit a particular symbiont genotype confers to its host. Partner quality not only impacts plant fitness, but also the nutritional composition of leaves as high-quality rhizobia fix more atmospheric N that plants incorporate into their tissues. Thus, results from GWAS can illuminate the G→P map of partner quality and generate novel candidate genes associated with variation in this ecologically and economically important trait.

Here we conduct multiple GWAS to study the complex genetic architecture and layers of context-dependency in the G→P map of partner quality. Because symbiotic extended phenotypes by definition have a multi-genomic basis (i.e., are influenced by multiple loci in both the host and symbiont), these traits are likely governed by many genomic variants, each with small effects. Results from a single GWAS may therefore be under-powered to identify the causal loci governing these traits. By examining associations that overlap across multiple GWAS using different host and symbiont genotypes, we can identify candidate loci that are consistently associated with traits regardless of environmental or genetic contexts, and thus, are good targets for subsequent functional validation, genetic engineering and coevolutionary studies. Using data from four separate GWAS that paired two lines of the host *Medicago truncatula* with 191 strains of the model rhizobium *Sinorhizobium meliloti*, we ask whether the genetic architecture of rhizobium partner quality differs across environments and across host genotypes (i.e., associations are “conditional”, governed by G × E and/or G × G), or alternatively, if variation at a core set of rhizobium genes control partner quality in all contexts (i.e., associations are “universal”, consistent in direction and magnitude across experiments and host genotypes). If the underlying genes that contribute to variation in partner quality are largely conditional, then predicting how partner quality evolves under different conditions will be more challenging. Additionally, we examine how the three genomic elements of *S. meliloti* contribute to partner quality variation across two host genotypes and in multiple environments, validating the functional division of labour of these elements in nature. Although the canonical symbiosis genes located on the symbiosis plasmids are likely to contribute, additional loci on the chromosome may be just as important for determining quantitative variation in partner quality. Finally, we suggest candidate metabolic functions and genetic loci that are responsible for both context-dependency as well as universally-beneficial effects that are consistent across environments.

## RESULTS

Using a panel of 191 strains of the model rhizobium *Sinorhizobium meliloti* isolated from natural populations in the native range, we conducted four separate GWAS that involved inoculating plants individually with each strain across two “replicate” experiments for two host lines of *Medicago truncatula:* for experiments 1 and 3, we used host line DZA (planted in March and September, respectively), while for experiments 2 and 4, we used A17 (planted May and November, respectively). Conditions between replicate experiments were kept as consistent as possible, the only major difference being the time of year they were planted. Our phenotypic and genetic analyses for these four experiments suggest important roles for context-dependency in the genetic control of symbiotic extended phenotypes. Based on our permutation method to determine significance, we found a total of 1,453 variants (**Supp. Dataset S1**, 67) in 746 (**Supp. Dataset S2**, 67) unique coding genes associated with shoot biomass, our focal rhizobium partner quality phenotype, while 5,402 variants (**Supp. Dataset S3**, 67) and 1,770 coding genes (**Supp. Dataset S4**, 67) were associated with at least one partner quality phenotype analyzed in this study (i.e., shoot biomass, leaf chlorophyll A content, plant height, leaf number); an additional 572 variants fell into 369 distinct non-coding regions (**Supp. Dataset S3**, 67). However, consistent with an *a priori* expectation that symbiotic extended phenotypes are governed by many variants with small effects, we indeed found that within a single GWAS, variant effect sizes tended to be small, with few approaching significance after controlling for the family-wise error rate (**Supp. Fig. S1**). Thus, the causal variants underlying partner quality are unlikely to be determined from a single GWAS, highlighting the need for multiple GWAS or other corroborating evidence before pursuing functional validation of particular candidate loci.

Our permutation approach nonetheless allows us to characterize the genetic architecture of partner quality by quantifying the number and location of associations across experiments and host genotypes, and thus, the degree to which these associations are conditional or universal. Three categories of G→P associations emerged from our analyses (**Fig. 1**; see **Supp. Fig. S2** in 68 for other traits). First were conditional associations that depended on the experiment (G × E: comparing experiments within host lines, **Fig. 1A**); 510 (68%) were found in only one of the four experiments. Second were “G × G genes”, the 51 (7%) genes that were associated with partner quality in both experiments with one host genotype but were found in neither experiment with the other host genotype (**Fig. 1A**). Specifically, 18 A17-specific genes were found in both experiments with this host (but never with DZA), while 33 DZA-specific genes were found in both experiments with this host (but never with A17). Last, but not least, were “universal” genes, the 60 (8%) genes that were found to be associated with rhizobium partner quality independent of host genotype or experiment (union in the center of **Fig. 1A**). Specifically, while only five of these genes were associated with partner quality in all four experiments, 55 additional genes were found in three of the four experiments and thus are strong candidates for contributing to partner quality in many conditions. We discuss these three types of genetic effects in turn, at the phenotypic, genomic (three elements in the tripartite genome), gene, and variant level, then conclude with the general implications of our findings for plant-microbe interactions and symbiosis evolution more broadly.

**FIG 1.**
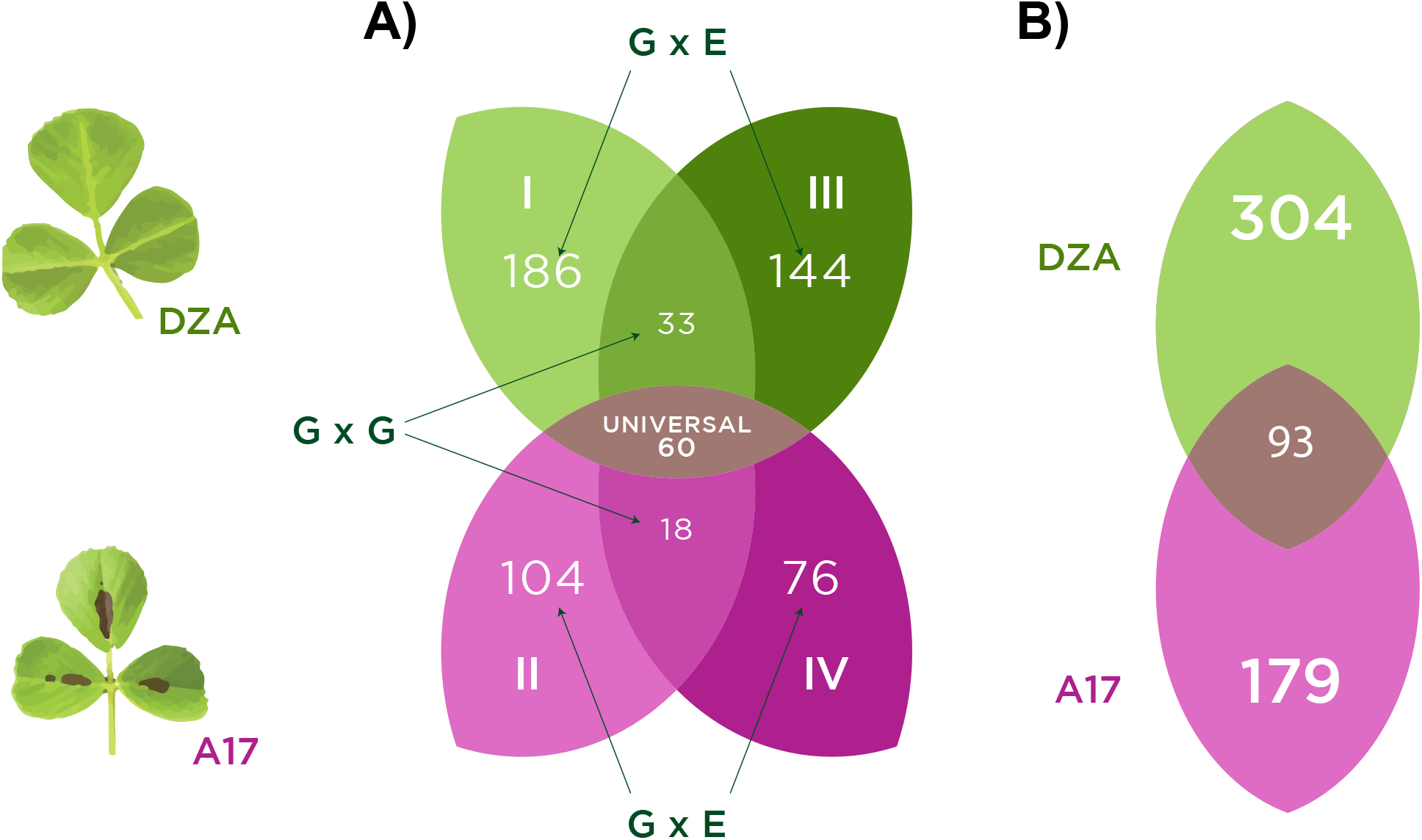
Association mapping of partner quality reveals strong signatures of environmental-dependence. Venn diagram showing number of rhizobium (*S. meliloti*) genes significantly associated with plant (*M. truncatula*) aboveground biomass (one metric of symbiotic partner quality) in **A)** each of four separate mapping experiments with either host line DZA (I and III in green) or host line A17 (II and IV in pink). Green and pink unions represent genes contributing to plant biomass in both experiments with either DZA or A17, respectively, while the mauve oval in the center represents universal genes found to contribute to plant aboveground biomass in at least three of the four experiments. **B)** Genes significantly associated with cross-experiment plasticity in plant aboveground biomass for host genotype DZA in green (experiments I vs. III), A17 in pink (experiments II vs. IV), or independently associated with plasticity in both hosts (central mauve triangle).

### Genotype-by-environment interactions in the G→P map of symbiotic partner quality

Because our different “environments” are actually replicate experiments, our goal here was to capture how reproducible associations are across experiments, rather than to track the specific environmental differences that contribute to conditional associations. We found that environmental dependence was a pervasive pattern at both the phenotypic, genomic, and individual gene levels, suggesting that different sets of rhizobium genes contribute to symbiotic partner quality under different environmental conditions. At the phenotypic level, ANOVA indicated abundant genetic variance in partner quality traits, including both significant main effects of strain as well as strong strain-by-experiment (G × E) interactions (see **Table 1** and within-experiment heritabilities in **Table 2**). Cross-experiment genetic correlations (*r*) of strain means between experiments within a host line were generally significant (**Table 2**), consistent with strain main effects (some strains had consistently higher partner quality than others; see **Supp. Figs. S3 & S4** in 68 for all within and between experiment trait correlations). Nevertheless, for most traits in both hosts, G × E was driven by changes in the rank order of strain means across experiments (**Table 2** crossing (%); see **Supp. Fig. S5** in 68 for reaction norms). Rhizobium strains varied considerably in cross-experiment plasticity for shoot biomass, with some strains having consistently low/high partner quality (i.e., resulting in small/large plant biomass, respectively) independent of the experiment, while others responded strongly to the environmental differences between experiments (**Supp. Fig. S6**, 68).

**TABLE 1.** Linear mixed model genotype by environment (G × E) ANOVAs for square-root transformed traits measured on plant lines A17 (left) and DZA (right). Numbers outside of and within parantheses in columns “A17” and “DZA” represent *χ*^2^ values and degrees of freedom. Rack included as a random effect, while all other terms are fixed. Significance of rack determined by calculating the log likelihood ratio between models with and without the random effect of rack.

| Trait | Model term | A17 | DZA |
| --- | --- | --- | --- |
| Chlorophyll | Intercept | 619.4(1)*** | 1447.65(1)*** |
|  | Strain | 252.6(190)** | 361.67(190)*** |
|  | Experiment | 0.15(1) | 0(1) |
|  | Strain X Experiment | 191.38(182) | 205.16(183) |
|  | Rack | 30.86(1)*** | 23.91(1)*** |
| Plant height | Intercept | 373.58(1)*** | 355.55(1)*** |
|  | Strain | 340.68(190)*** | 268.05(190)*** |
|  | Experiment | 20.24(1)*** | 2.5(1) |
|  | Strain X Experiment | 308.63(185)*** | 177.71(183) |
|  | Rack | 38.35(1)*** | 54.12(1)*** |
| Leaves | Intercept | 207.69(1)*** | 164.09(1)*** |
|  | Strain | 355.37(190)*** | 207.58(190) |
|  | Experiment | 7.85(1)** | 0.45(1) |
|  | Strain X Experiment | 243.88(185)** | 218.54(183)* |
|  | Rack | 27.51(1)*** | 41.67(1)*** |
| Shoot biomass | Intercept | 97.65(1)*** | 179.32(1)*** |
|  | Strain | 544.81(190)*** | 499.67(190)*** |
|  | Experiment | 10.08(1)** | 6.93(1)** |
|  | Strain X Experiment | 330.06(185)*** | 262.2(183)*** |
|  | Rack | 11.32(1)*** | 155.67(1)*** |

Significance: $p < 0.001 = \text{'***'}$ ; $p < 0.01 = \text{'**'}$ ; $p < 0.05 = \text{'*'}$

**TABLE 2.** Within-experiment broad-sense heritabilities (H^2^) and cross-experiment genetic correlations (*r*). G × E interactions were partitioned into changes in variance versus rank order (i.e., crossing), and the percent due to crossing is presented. Experiments (exps.) 1 and 3 for DZA, 2 and 4 for A17.

| Trait | Plant line | $H^2$ (exps. 1 or 2) | $H^2$ (exps. 3 or 4) | $r$ | Crossing (%) |
| --- | --- | --- | --- | --- | --- |
|  | DZA | Exp. 1 | Exp. 3 |  |  |
| Shoot biomass |  | 0.248*** | 0.253*** | 0.265*** | 87.19 |
| Chlorophyll |  | 0.118** | 0.141*** | 0.297*** | 99.06 |
| Plant height |  | 0.131** | 0.065* | 0.175* | 99.97 |
| Leaves |  | 0.05 | 0.084** | -0.071 | 63.96 |
|  | A17 | Exp. 2 | Exp. 4 |  |  |
| Shoot biomass |  | 0.303*** | 0.262*** | 0.221** | 99.83 |
| Chlorophyll |  | 0 | 0.093** | 0.201** | 0 |
| Plant height |  | 0.181*** | 0.173*** | 0.1 | 73.37 |
| Leaves |  | 0.173*** | 0.127*** | 0.173* | 94.13 |

Significance: $p < 0.001 = \text{'***'}$ ; $p < 0.01 = \text{'**'}$ ; $p < 0.05 = \text{'*'}$

At the genomic-level, our evidence indicates that G × E was driven by small-scale shifts in the identity, and estimated allelic effects, of individual loci across experiments rather than large shifts in which genomic elements have contributed to partner quality variation (**Supp. Fig. S7**, 68). The vast majority of genes contributing to partner quality variation were located on the symbiotic elements (megaplasmid pSymA or chromid pSymB; **Fig. 2**; **Supp. Dataset S2, 67**; see **Supp. Fig. S8** in 68 for other traits), although some environmentally-dependent loci were found on the chromosome (i.e., G × E and plasticity). Within host lines, 330 total genes were mapped with host line DZA (186 and 144 in experiments 1 and 3, respectively), while only 180 total genes mapped to host line A17 (104 and 76 in experiments 2 and 4, respectively; **Fig. 1A**).

**FIG 2.**
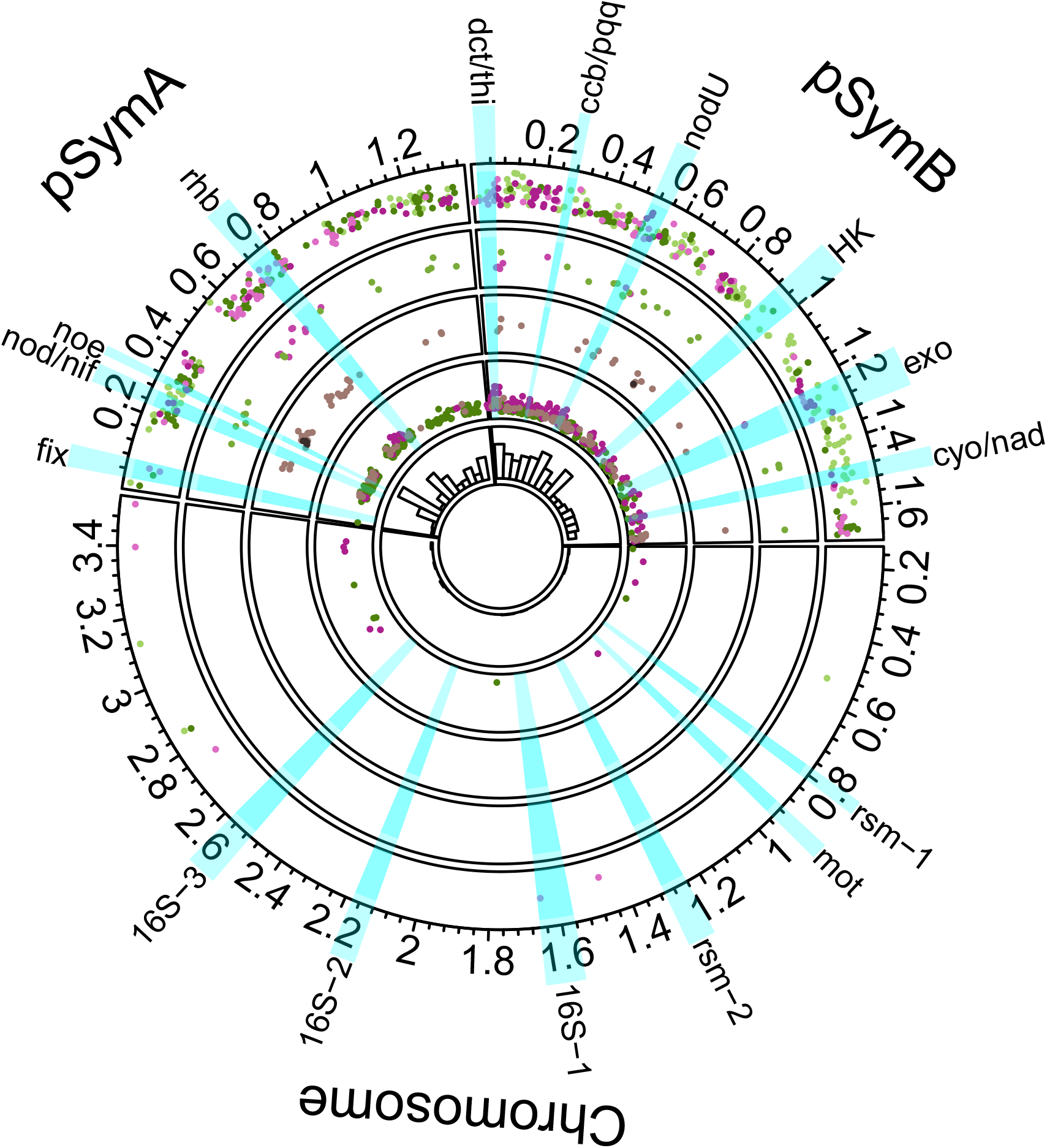
Loci associated with partner quality are mostly limited to the symbiosis plasmids. Circos plot showing positions of genes (dots) significantly associated with shoot biomass. Each ring represents a different gene category, outermost to innermost: 1) G × E, 2) G × G, 3) partially universal and universal, 4) plasticity, while 5) depicts a histogram based on the total number of significant genes across 100 kbp-sized windows. The x- and y-axes for rings 1-4 represent genomic position (Mbp) and average absolute effect sizes of variants within each gene, respectively. The colours reflect categories in the Venn Diagrams: for rings 1, 2, and 4, genes associated with DZA-only traits are represented by shades of green, on A17-only with shades of purple, and both hosts in mauve (ring 4). For ring 3, genes associated with both hosts in more than three environments are represented in mauve (i.e., “partially universal”), and universal genes in black. Relevant loci are highlighted in blue, with abbreviations for clusters on the outer circle as specified in **Supp. Fig. S8** (67).

At the gene-level, for experiment 1 in host line DZA, the INTERPRO terms “Transcription regulator *hth, lac*I and “Lambda repressor-like, DNA-binding domain” were marginally significantly enriched (p = 0.00101 and 0.00482, respectively), as was the “Oxidative phosphorylation” KEGG pathway (p = 0.00959; **Supp. Dataset S5**, 67), whereas for experiment 3 in host line DZA, the “Benzoate degradation” KEGG pathway was significantly enriched (FDR-corrected p = 0.00880), while the GO terms “3,4-dihydroxybenzoate catabolic process” and “transmembrane transport” were marginally significantly enriched (p = 0.00204 and 0.00376, respectively; **Supp. Dataset S5**, 67). For experiment 2 in host line A17, the GO terms “transcription, DNA-templated” were significantly enriched (FDR corrected p = 0.00143), as were the related UNIPROT keywords for “transcription regulation” (FDR corrected p = 0.00310) and “DNA-binding” (p = 0.00310; **Supp. Dataset S5**, 67), whereas for experiment 4 in host line A17, the INTERPRO term “Amidohydrolase 1” was marginally significantly enriched (p = 0.0307; **Supp. Dataset S5**, 67). Overall, many of the underlying molecular processes important for driving variation in partner quality appear to be environmentally-dependent, making it harder to predict which genes and pathways will be important for determining symbiotic outcomes under different conditions.

At the variant-level, for each host (DZA or A17), we used correlations of the estimated allelic effects to assess the degree to which the individual effects of rhizobium alleles on partner quality were consistent in direction and/or magnitude, or whether they depended on the experiment. When variants were significantly associated with partner quality in both experiments (**Fig. 3** dark points), they tended to have inconsistent effects on host line DZA (**Fig. 3A**), while being more consistent on A17 (d**Fig. 3B**). In fact, on A17, all of the nearly-universal variants (**Fig. 3B** black dots) had concordant (i.e., same-sign) effects on shoot biomass between experiments, whereas the opposite was true for DZA (**Fig. 3A** black dots) - all nearly-universal variants had discordant (i.e., opposite-sign) effects between experiments. For host line DZA, 12 particularly interesting variants had significant but opposing effects on plant biomass across the two experiments (**Fig. 3A**; **Supp. Dataset S1**, (67)). Such associations might point to interesting environmentally-dependent genes. Regardless of host line, however, the vast majority of variants in our studies had conditionally-neutral effects, even those with large magnitude (**Fig. 3**; see **Supp. Fig. S9** in 68 for other traits), and for both hosts, we rejected the global null hypothesis that allelic effects were the same across experiments (both p < 0.0001; **Supp. Fig. S10**, 68).

**FIG 3.**
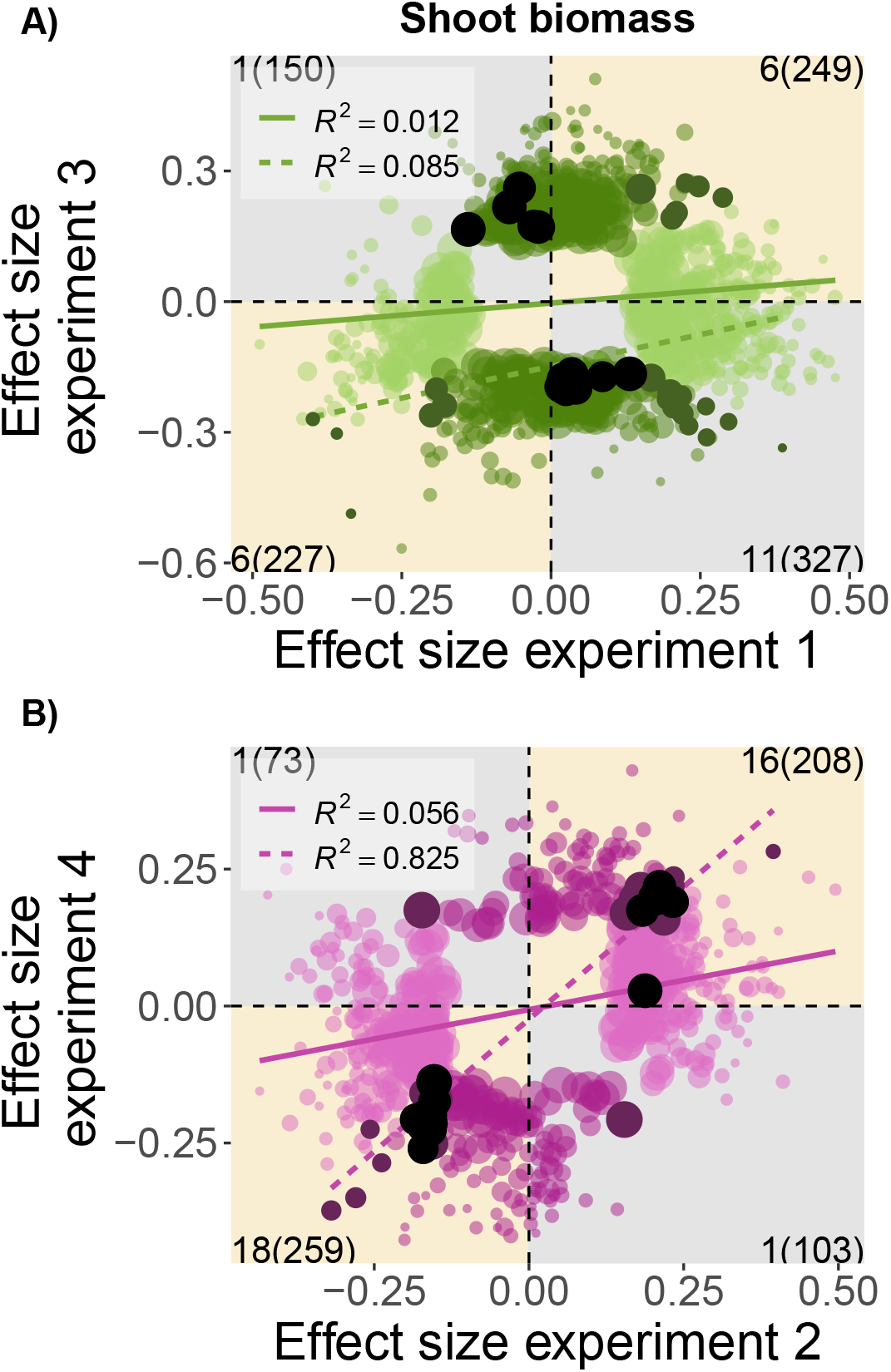
Allelic effects are more consistent between experiments for A17, but not for DZA. Correlations between the estimated allelic effects of individual *S. meliloti* variants on plant shoot biomass (from GWAS) in each of two experiments for either **A)** host DZA (green) or **B)** A17 (pink). Only allelic effects that were significant in one (lighter colours) or both (dark points) environments are shown, while black dots represent nearly universal variants, i.e., associated with the same trait in three experiments. Linear relationships and R^2^ values are depicted for all variants (solid coloured line) or variants significant in both experiments (dotted coloured line). Counts of significant variants for one or both environments appear in the corners of each quadrant within or outside the parentheses, respectively.

Next we mapped the among-strain differences in plasticity (**Supp. Fig. S6**, 68) to identify specific loci contributing to G × E. At the genomic-level, most (63%) plasticity loci were found on pSymB (**Fig. 2**; **Supp. Fig. 7B**, 68). Of 576 genes, 400 (69%) were also associated with shoot biomass variation *within* at least one of the four experiments when each was mapped independently, indicating abundant overlap in the genetic architecture of both within- and among-experiment partner quality variation (**Supp. Dataset S6**, 67). Plasticity loci were marginally significantly enriched for the INTERPRO terms “Ti-type conjugative transfer relaxase *tra*A” (p = 0.0292) and “*mob*A/*mob*L protein” (p = 0.0292), as well as several INTROPRO terms containing “tetratricopeptide-repeat” (p range: 0.006 to 0.0310), among others (**Supp. Dataset S6**, 67). Of 576 total plasticity genes, only 93 (16%) were mapped in both host genotypes (**Fig. 1B, Fig. 2**; **Supp. Dataset S7**, 67). Perhaps most interesting are the 14 of these plasticity genes that were *not* associated with variation within any of the four experiments when mapped separately (**Supp. Dataset S6**, 67); these loci are particularly strong candidates for understanding how G × E in rhizobia scales up to alter host growth, and none to our knowledge have known functional roles in symbiosis.

### The genetic architecture of rhizobium partner quality depends on host genotype

Given our four-experiment design, we used the conservative approach of only considering G × G genes to be those that were associated with partner quality in *both* experiments with one host genotype and *neither* of the other. We found 33 genes that contributed to partner quality but only in DZA, and 18 that contributed but only in A17 (**Fig. 1**; **Supp. Dataset S2**, 67). These genes split evenly across the two symbiosis plasmids of the genome (pSymA: N = 24; pSymB: N = 27). Below, we discuss our results at the gene-level for each host line separately.

In the list of A17-only G × G genes, analyses of UNIPROT keywords indicated that “Selenium” was marginally significantly overrepresented (p = 0.00771), while GO terms for “transferase activity, transferring glycosyl groups” were marginally over-represented (p = 0.0417; **Supp. Dataset S5**, 67). Notably, these genes include both *exo*U (SMb20948/ NP_437611.1) and *exo*W (SMb21690/NP_437613.1; **Supp. Dataset S2**, 67), two glucosyltransferases critical for adding the 6^th^ and 7^th^ sugars (respectively) on the succinoglycan molecule, an exopolysaccharide required for infection thread formation and thus host plant invasion for effective symbiosis (69, 70, 71). Non-functional *exo* genes are known to result in less-efficient, though sometimes not entirely deficient, symbiosis (72). Together with this prior knowledge, our finding that variants in two succinoglycan biosynthesis enzymes affected host biomass production in host A17 suggests a strong role for succinoglycan-mediated host invasion in determining symbiotic benefits in natural populations. Future studies of coevolution between succinoglycan structure and host plant detection would likely be fruitful. Also notable is *sel*A(SMa0011/NP_435251.1; **Supp. Dataset S2**, 67), a transferase required for the biosynthesis of selenocysteine, a less-used amino acid incorporated into select proteins in only about 20% of bacterial genomes (73, 74, 75). To our knowledge, nothing is known about the role of selenocysteine in rhizobia or in legume-rhizobium symbiosis.

By contrast, the list of DZA-only G × G genes was dominated by GO terms “transcription factor activity, sequence-specific DNA binding” (p = 0.0247) and “cellular amino acid metabolic process” (p = 0.0425; **Supp. Dataset S5**, 67). Notably this list contains three *lys*R transcriptional regulators (**Supp. Dataset S2**, 67) of at least 90 in the genome: SMa2287 (NP_436477.2) on pSymA plus SM_b20494 (NP_437016.1) and SM_b21434 (NP_437803.1) on pSymB are not well-studied, and had weak (SM_b20494) or undetectable effects in a plasmid insertion mutagenesis screen compared to the named *lsr*A and *lsr*B (76). This class of loci is well-known to control expression of genes for symbiosis, both mutualistic and pathogenic (77). Beyond those discovered in knockdown studies, our results suggest additional symbiotic roles for natural variation at these *lys*R regulators in *S. meliloti*. Interestingly, one metabolic process gene (*gdh*A: SMa0228/NP_435368.1 on pSymA; **Supp. Dataset S2**, 67), part of one bacterial pathway for assimilation of ammonium into glutamate (78), was identified in a comparative genomics study of five *Sinorhizobium* (formerly *Sinorhizobium*) species as specific to *S. meliloti* (79). The role of *gdh*A in rhizobial symbiosis is not well-known, though interestingly it was found to increase ammonium assimilation when the *E. coli* copy was expressed transgenically in tobacco (80).

### Universal associations highlight transport functions and secretion systems

Despite these layers of context-dependency, we did find significant main effects of strain for all phenotypes (**Table 1**). Concomitantly we resolved a number of loci that were consistently associated with shoot biomass, being mapped in either three experiments (55 nearly-universal genes) or even all four experiments (5 universal genes) (**Fig. 1**). At the genomic and gene-levels, the set of 60 universal/nearly-universal genes were split between pSymA and pSymB (38 and 22 loci, respectively; **Fig. 2** mauve dots; **Supp. Dataset S2**, 67) and featured marginally-enriched INTERPRO terms for “Tetratricopeptide-like helical” (p < 0.001) and “Tetratricopeptide repeat-containing domain” (p = 0.00160) as well as UNIPROT keywords “Transmembrane” (p = 0.0190) and “Transmembrane helix” (p = 0.0407), though none of these terms were significant after FDR-correction (**Supp. Dataset S5**, 67).

Of the five truly universal genes (mapped in all 4 experiments; **Supp. Dataset S2**, 67), most do not have known functions in symbiosis. This includes a *cax* gene (SMa0675/ NP_435603.2) putatively involved in calcium/proton exchange. Though *cax* genes are widespread in bacteria and eukaryotes (81), and the importance of calcium both in nodule establishment and trade of benefits is known (82, 83), it is difficult to hypothesize on the function of this particular gene or its genetic variants in symbiosis currently. Given their as-yet unknown functions and lack of context-dependency in our studies, the five universal candidates might hold the most potential for novel functional information and consistent phenotypic effects, which might make them ideal candidates both for validation and for symbiosis improvement.

Like the five universal genes, the “nearly-universal” set of genes associated with shoot biomass in three (of four) experiments highlights the existence of segregating natural variation in several interesting metabolic pathways, only some of which have established roles in symbiosis. We found five loci annotated as involved in transmembrane transport (**Supp. Dataset S2**, 67), including *kdp*A (SMa2333/NP_436501.1), *pot*E (SMa0678/NP_435605.1), and *msb*A1 (SM_b20813/NP_437093.2) - in addition to the nodulation protein *glm*S/*nod*M (SMa0878/NP_435728.1) and the universally-associated *cax* transporter (discussed above). The role of potassium transporters such as *kdp*A in osmoregulation during symbiosis is not well understood (84). The *pot*E locus codes for a putrescine/ornithine antiporter, while another nearby nearly-universal locus (SMa0682/NP_435607.4) is a predicted amino acid decarboxylase in the putrescine biosynthesis pathway, potentially suggesting a role for variation in putrescine metabolism, which is known to vary in *S. meliloti* (45), in symbiotic partner quality (45, 85).

Finally, we interrogated the gene sets from two key studies that have associated natural variation in *S. meliloti* genomes with symbiotic partner quality (see **Supp. Dataset S2**, “overlap” column; 45, 66). Our nearly-universal gene set contained eight loci that overlapped with the top 100 associations with A17 biomass from Epstein et al. (45). Most notable is the fructose-6-phosphate aminotransferase *nod*M/*glm*S (SMa0878/NP_435728.1) that catalyzes a precursor of both peptidoglycan and Nod factor in the glucosamine biosynthesis pathway. This locus is located in the symbiosis gene region of pSymA, though a paralog exists on the chromosome (SMc00231/NP_385762.1; 86). Knockout mutants of *nod*M are known to decrease N-fixation of *S. meliloti* on alfalfa (87) and *Rhizobium leguminosarum* (88); together with Epstein et al. (45), our studies highlight the role of natural variation in bacterial glucosamine metabolism in determining plant health. We also found six genes in this nearly-universal set that were also associated with symbiotic partner quality, rhizobium fitness, or both in the experimental evolution study of Batstone et al. (66). Most notable are two *tra* (transfer) loci (*tra*A2 on pSymB and *tra*G on pSymA), potentially part of a Type IV Secretion System (T4SS) responsible for targeting proteins to host cells (89, 90). While the variants we found in these loci are segregating in natural populations in the native range of *S. meliloti*, these loci also evolved *de novo* in response to passaging through the same host for multiple generations (66), making them strong candidates for a consistent role in symbiosis.

## DISCUSSION

Understanding our symbiotic world requires a genetically-accurate appreciation of the symbiotic extended phenotypes upon which selection acts. While evolutionary ecology has long recognized the importance of genetic variation in symbiosis while ignoring the underlying mechanisms, functional geneticists have traditionally resolved mechanisms without taking into account natural variation. Synthesis of these two perspectives has started to resolve ecologically-relevant quantitative variation at the nucleotide level (45, 46, 47, 60, 91, 92). Here we quantify multiple symbiotic extended phenotypes in four GWAS using a model plant-microbe symbiosis and find that the genetic architecture of symbiotic partner quality is complex, underlain by networks of numerous interacting loci and environmental-dependence. We find that some loci in the microbial symbiont have consistent effects on host growth across experiments, contributing to the overall differences in mutualistic partner quality that have been the focus of many empirical and theoretical studies of mutualism to date (reviewed by 40, 93, 94, 95, 96). Nonetheless, most loci identified in our study were significantly associated with variation in partner quality in specific environments (G × E effects), or with specific host genotypes (G × G effects). We first discuss the roles of environmentally-dependent loci, versus universal loci that are found consistently. Next we discuss the coevolutionary implications of genotype-dependence (G × G interactions), then wrap up with a call for further synthesis with metabolic network models towards systems genetics of symbiosis.

### Environmental context-dependency in the G→P map of symbiosis

Ecological effects of context-dependency in mutualism have been recognized for a long time, i.e., traits and mutualism benefits often shift across environmental conditions such as nutrient availability or light environments (97, 98, 99), although not ubiquitously for all traits (e.g., 100, 101). More recent studies have begun to document the evolutionary changes that can result from these ecological effects, e.g., divergence of host and/or symbiont symbiosis traits across strong ecological gradients (59, 102, 103, 104, 105). Evolutionary change in response to environments implies that the loci underlying selected traits have differential effects on fitness across environments. Here we identify the loci that generate important trait variation both within and among environments, the trait variation upon which selection acts in nature.

The majority of context-dependent partner quality genes we identified might be viewed much like the conditionally-neutral variation so often found in studies of local adaptation (106, 107, 108, 109, 110), contributing significantly to variation in partner quality in some contexts but having a range of weaker effects (both in the same or opposing direction) in another context. For example, by experimentally-evolving strains of *S. meliloti* on five *M. truncatula* lines, Batstone et al. (66) found that local adaptation was largely governed by conditional neutrality (beneficial on local host, neutral on non-local hosts) or mildly deleterious effects on non-local hosts, likely due to drift in the local context. Yet, in nature where rhizobium population sizes are much larger and more diverse, and gene flow is present, the extent to which local adaptation occurs has rarely been tested but might be unlikely (28, 111, 112), except for populations differentiated by strong ecological gradients (e.g., 102). Moreover, the host genotypes used in our study, DZA and A17, are unlikely to share an evolutionary history with our strains, and so it remains unclear whether stronger trade-offs (and less conditional neutrality) would be present if our strains shared an evolutionary history with the host lines being tested.

Despite widespread conditional neutrality, a handful of interesting variants had strong effects on host growth, but in *opposing directions* across experiments within a single host genetic background. These sorts of antagonistic effects can favour different variants in different environments, and thus, potentially help explain the maintenance of mutualism variation in nature (95, 113, 114, 115). Nevertheless we note that these sorts of G × E variants were rare in our study, despite strong rank-order effects among strain means at the organismal level (**Table 2**); moreover, because our “environments” were simply different greenhouse experiments, relating such antagonistic effects to adaptation in the wild will require the type of *in situ* studies that are common in plants (e.g., 104, 107, 116) but more difficult in soil microbes.

We nevertheless found several genes and pathways that are ideal for functional follow-up studies aimed at identifying consistent associations, or loci which might benefit from “fine-tuning” symbiotic benefits towards improving plant health. The strengths of GWAS (capturing genetic variation, ecological relevance) also lead to weaknesses (imprecision due to confounding population structure, false negatives and false positives). Thus our strongest recommendations for functional hypotheses for validation in follow up studies are the universal genes identified here that overlap with candidate genes identified in separate GWAS using different genomic backgrounds and environments (e.g., 45, 66).

Overall we find that the loci underlying quantitative variation in symbiotic extended phenotypes often (but not always) depend on the environment, and therefore that a nuanced understanding of how complex traits interact with environmental variables will be necessary for many of the lofty goals in plant microbiome and symbiosis research (9, 35). This point has been made before (117), as the presence of G × E as studied at the phenotypic level has been recognized for decades; what is novel here is our ability to interrogate this variation at the genomic level and for multiple host lines (see below).

### Coevolutionary implications of genotype-dependence

Uncovering the complex genetics of how two (or more) genomes interact with each other to generate trait variation is an important step to better understanding how these traits (co)evolve (6) and how to better manipulate traits in the future to address societal challenges (37, 118). For example, identifying the loci underlying mutualistic traits allows us to address longstanding debates within mutualism theory (119), including how readily conflict evolves (120), and how genetic variation is maintained despite host selection for the ‘best’ symbiont (95).

Previous mapping efforts in the legume-rhizobium system have focused on the *Medicago* HapMap collections, which maximize host diversity using a range-wide sample. Our study focuses on the segregating natural variation within a symbiont at a smaller geographic scale, a scale at which pSymA and pSymB segregate (112); thus the G × G-driven variation we find here would be available to local evolutionary and/or coevolutionary processes. G × G interactions for fitness outcomes have long been of interest in the legume-rhizobium symbiosis (25, 121, 28, 102, 122) and other interactions (123, 124, 125) because such statistical interactions generate the fitness variation that drives coevolution. Additionally, G × G interactions have implications for breeding and agricultural production because the functional effects of symbiont variation are likely to depend on the crop genotype (126, 127). Recent molecular genetic and transcriptomic approaches on a handful of genotypes have begun to resolve the mechanistic underpinnings of G × G (128, 129, 130, 131, 132, 133), while biparental or GWAS mapping approaches (45, 46, 134) provide broader insight into the genetic architecture underlying G × G (i.e., the number, average effect size, and consistency of loci). The picture emerging from ours and others’ studies is that G × G, like symbiotic partner quality itself, has a complex, polygenic basis and will require both statistically sophisticated and metabolically-informed models to unravel.

### Symbiotic extended phenotypes as quantitative traits

In the age of rapid microbiome sequencing and expanding efforts to characterize the loci in plant genomes that contribute to microbiome variation among cultivars or genotypes (e.g., rice, *Lotus, Medicago*), our results present an important juxtaposition, as the abundant and context-dependent genetic variation characterized in detail here occurs within a set of 191 rhizobia strains with > 98% average nucleotide identity (well above the typical threshold for delineating and enumerating operational taxonomic units, or OTUs, in metagenomic studies). At the same time, functional genetic studies have made much progress identifying loci critical to symbiosis establishment and downstream processes by creating knock-out or knock-down mutants and comparing their associated symbiotic phenotypes to a wildtype strain (e.g., *dnf* mutants in legumes, 135; fix+/fix-mutants of rhizobia, 136), but these loci are often viewed as on-or-off switches for symbiosis more generally, or cooperation more specifically in models of mutualism theory (137, 138, 139, 140). Our study demonstrates that most loci within the symbiont genome act more like dials than on-and-off switches, generating the quantitative variation in symbiotic extended phenotypes observed in nature.

While our approach allows us to generate novel candidate loci that are consistently associated with partner quality across contexts, *in silico* modelling of both plant (141) and rhizobium (142) metabolism that links together genetic information with metabolic pathways could then be used to simulate key symbiotic processes under a wide range of conditions (143). Ultimately combining transcriptomic, genomic, and metabolomic datasets will be required for a synthetic understanding of how nucleotide variation percolates up through shared symbiotic metabolic networks (17, 144, 145). Efforts to reintegrate research on symbiosis genetics and (co)evolution with plant-microbiome work (reviewed in 37, 118) will be fruitful in revealing additional intraspecific variation driving patterns of genotype-dependence and coevolution and resolving mechanisms of host control of the microbiome.

## MATERIALS AND METHODS

Full details are available in the Supplemental Materials (**Supp. Methods**, 68). We performed four greenhouse experiments to estimate partner quality phenotypes in *S. meliloti*. In each experiment, plants from one of two host lines (either A17 or DZA) were grown in single inoculation with each of 191 *S. meliloti* strains, with three to four replicates per strain per experiment (six to eight total replicates for each plant line × strain combination, N = 2,825 plants total). Experiments were planted in 2018, in March (I: DZA), May (II: A17), Sept (III: DZA), and Nov(IV: A17). Uninoculated controls (40 per experiment) were included to gauge contamination, which was minimal overall, and limited to the first two experiments (**Supp. Fig. S11**, 68). We measured multiple proxies of partner quality, namely leaf chlorophyll A content, plant height, number of leaves, and above-ground dried shoot biomass, although we focus on the latter in the main text (see Supplemental Materials for all others). We conducted multiple phenotypic analyses to determine how much variation in partner quality was due to strain, experiment, and the interaction of both, among other questions.

We sequenced the entire genomes of all 191 *S. meliloti* strains, called single nucleotide polymorphisms (SNPs, henceforth referred to as variants), and performed four separate GWAS that accounted for rhizobium population structure and included only unlinked variants. We determined which variants were significantly associated with partner quality using a permutation method (45) that involved generating 1000 randomized datasets (i.e., genotypes randomized with respect to phenotypes) and running a linear mixed model (LMM) on each. In all models, we included the same set of variants as well as the kinship matrix as a random effect to account for associations that arise due to population structure. However, because we conducted our experiments under controlled greenhouse conditions, unmeasured variables that are population-stratified are unlikely to confound our results. We then tagged variants from the non-randomized run that fell above the 95% false discovery rate cut off based on the combined randomized runs. Although more computationally demanding and less conservative than conventional Bonferroni correction, the advantage of our permutation approach is that it better captures the unique properties specific to each dataset such as trait distributions, patterns of linkage disequilibrium, and missing data, while also controlling for the per-variant false positive rate in the presence of associations at other loci. Thus, despite the inherent challenges associated with determining the causal variants underlying highly polygenic traits (i.e., numerous variants with small effects), our permutation approach nonetheless allows us to characterize the genetic architecture of partner quality, determine the degree to which associations are conditional or universal, and even identify candidate loci that are most likely to contribute to variation in partner quality across conditions by comparing the variants we identified as “universal” with those highlighted in other GWAS using different experimental conditions and host and symbiont genotypes.

Based on our permutation method, we binned the resulting significant variants into three categories based on the context-dependency of their phenotypic effects, and thus, their contribution to the layers of the G→P map for each of our symbiotic extended phenotypes. First, “nearly-universal genes” were those found to have significant effects in at least three of the four experiments for a particular trait (“universal genes” were mapped in all four experiments). Second, we used a conservative approach to call “G × G genes” as those mapped in both experiments for one host genotype but neither of the experiments for the other host genotype (i.e., “DZA G × G” genes were significant in both experiments I and III with DZA but neither II nor IV with host A17). Third were genes significantly associated with partner quality in a single experiment, and never in another (i.e., “G × E” genes). Finally, we conducted candidate gene functional analyses to understand how loci within different categories differed from one another functionally, or whether they were part of the same networks/metabolic pathways.

### Data Availability

Strains and plant lines are available upon request. All raw data and analysis code are available on GitHub (see “Complex_genetics” folder). Once raw sequence reads and assemblies are archived and made available on NCBI, accession numbers will be added to this manuscript.

## SUPPLEMENTAL MATERIAL

Detailed methods and thirteen (13) supplementary figures are available on Zenodo (68) (doi: https://doi.org/10.5281/zenodo.5550958). Seven (7) supplementary datasets are available on Dryad (68) (doi: https://doi.org/10.5061/dryad.5dv41ns6r). All supplemental material, in addition to the main text, is also available at bioRχiv(doi: https://doi.org/10.1101/2021.08.03.454976).

## ACKNOWLEDGMENTS

We thank greenhouse staff (esp. Debbie Black). We acknowledge funding from the National Science Foundation (IOS-1645875 and NPGI-1401864), a Carl R. Woese Institute for Genomic Biology postdoctoral fellowship to R.T. Batstone, strain sequencing by Joint Genome Institute (CSP-1223795), as well as analytical advice from B. Epstein.

