## Supplementary Materials for "Genome-wide association studies across environmental and genetic contexts reveal complex genetic architecture of symbiotic extended phenotypes"

Zenodo: https://doi.org/10.5281/zenodo.6712084 Zenodo: https://doi.org/10.5281/zenodo.6712084 Zenodo: https://doi.org/10.5281/zenodo.6712084 Zenodo: https://doi.org/10.5281/zenodo.6712084

#### SUPPLEMENTAL MATERIALS

### Genome-wide association studies across environmental and genetic contexts reveal complex genetic architecture of symbiotic extended phenotypes

**Rebecca T. Batstone,[a,*] Hanna Lindgren,[b] Cassandra M. Allsup,[b] Laura A. Goralka,[b] Alex B. Riley,[b] Michael A. Grillo,[c] Amy Marshall-Colon,[a,b] Katy D. Heath,[a,b,*]**

Carl R. Woese Institute for Genomic Biology, University of Illinois at Urbana-Champaign, 1206 West Gregory Drive, Urbana, IL 61801 USA[a]; Department of Plant Biology, University of Illinois at Urbana-Champaign, 286 Morrill Hall, 505 South Goodwin Avenue, Urbana, IL 61801 USA[b]; Department of Biology, Loyola University Chicago, 1032 W. Sheridan Road, Chicago, IL 60618 USA[c]

R. Batstone and H. Lindgren contributed equally. Author order was determined based on overall contribution, particularly time spent analyzing the data, as well as writing and compiling feedback.

**ABSTRACT** A goal of modern biology is to develop the genotype-phenotype (G→P) map, a predictive understanding of how genomic information generates trait variation that forms the basis of both natural and managed communities. As microbiome research advances, however, it has become clear that many of these traits are symbiotic extended phenotypes, being governed by genetic variation encoded not only by the host's own genome, but also by the genomes of myriad cryptic symbionts. Building a reliable G→P map therefore requires accounting for the multitude of interacting genes and even genomes involved in symbiosis. Here we use naturally-occurring genetic variation in 191 strains of the model microbial symbiont *Sinorhizobium meliloti* paired with two genotypes of the host *Medicago truncatula* in four genome-wide association studies to study the genomic architecture of a key symbiotic extended phenotype – partner quality, or the fitness benefit conferred to a host by a particular symbiont genotype, within and across environmental contexts and host genotypes. We define three novel categories of loci in rhizobium genomes that must be accounted for if we want to build a reliable G→P map of partner quality; namely, 1) loci whose identities depend on the environment, 2) those that depend on the host genotype with which rhizobia interact, and 3) universal loci that are likely important in all or most environments.

**IMPORTANCE** Given the rapid rise of research on how microbiomes can be harnessed to improve host health, understanding the contribution of microbial genetic variation to host phenotypic variation is pressing, and will better enable us to predict the evolution of (and select more precisely for) symbiotic extended phenotypes that impact host health. We uncover extensive context-dependency in both the identity and functions of symbiont loci that control host growth, which makes predicting the genes and pathways important for determining symbiotic outcomes under different conditions more challenging. Despite this context-dependency, we also resolve a core set of universal loci that are likely important in all or most environments, and thus, serve as excellent targets both for genetic engineering and future coevolutionary studies of symbiosis.

**KEYWORDS:** GWAS, mapping, *Medicago truncatula*, *Sinorhizobium meliloti*, symbiosis, partner quality, rhizobium, G x E, G x G, genotype-phenotype map, G→P map, root nodule, nodulation, symbiotic nitrogen fixation

Zenodo: https://doi.org/10.5281/zenodo.6712084 Zenodo: https://doi.org/10.5281/zenodo.6712084 Zenodo: https://doi.org/10.5281/zenodo.6712084 Zenodo: https://doi.org/10.5281/zenodo.6712084 https://doi.org/10.5281/zenodo.6712084



**CONTENTS:**

Methods
Datasets S1 to S7
References
Figures S1 to S13

#### SUPP. METHODS

**Study system.** *Medicago truncatula* Gaertn. is an annual legume native to the Mediterranean region in Southern Europe and Northern Africa, and it also occurs in Asia. It lives in a symbiotic association with nitrogen-fixing bacteria in the genus *Sinorhizobium* and has been used as a model-organism to study symbiotic interactions. We used two lines of *Medicago truncatula* in this experiment: A17 (Australia) and DZA 315.16 (Algeria; hereafter abbreviated as DZA). These lines and have been leveraged in studies establishing the role of variable plant-encoded nodule-specific cysteine rich peptides (NCRs, specifically those encoded by *nfs*1 and *nfs*2) in governing the level of nitrogen fixed (Fix+, Fix–, or intermediate) in particular host and strain combinations (1, 2, 3, 4); these lines are included in the *Medicago* HapMap panel of re-sequenced GWAS lines and RIL parents (http://www.medicagohapmap.org). We used 191 strains of the rhizobium *S. meliloti* in this study, which were isolated from the nodules of *M. truncatula* plants grown in soils from 24 sites in Corsica, France and Spain, as part of a larger effort to understand coevolution in this legume-rhizobium mutualism (5, 6).

**Experimental design.** Seeds were scarified with either a razor blade or sandpaper, then sterilized by first rinsing with 95% EtOH followed by soaking in commercial bleach (6% hypochlorite) for 7 min, rinsed thoroughly with water, and imbibed in $ddH_2O$ overnight at 4°C in the dark. Seeds were then transplanted into sterilized 107 ml SC7 Cone-tainers (Stuewe & Sons Inc., Tangent, OR), each containing an autoclave-sterilized mixture of Turface MVP calcined clay (Profile Products LLC, Buffalo Grove, IL) and the UIUC greenhouse's root wash mix (1:1:1 soil–calcined clay–torpedo sand) in equal volumes, for a final mixture of 1:4:1 soil–calcined clay–torpedo sand. Pots were randomized into racks in the greenhouse, with 10 pots per rack, and racks were evenly arranged across three benches.

*S. meliloti* cultures were grown in liquid tryptone-yeast (TY) medium (7) for 18-20 hrs at 30°C. Before inoculation, the cell density of each culture was measured with NanoDrop 2000c spectrophotometer (Thermo Fisher Scientific) and adjusted to $\sim 10^6$ cells/ml ($OD_{600}$ = 0.1) by diluting the cultures with sterile TY medium, when necessary. Each plant was inoculated with 500 ml of liquid culture 10-12 days after seeds were planted, and the soil surface received a thin (~0.5 cm) layer of sterile sand after inoculation to minimize cross-contamination. Plants were misted (1/2" M NPT upright misting nozzle, Senninger Irrigation Inc., Clermont, FL) four times per day for 45 min at a time for the first two weeks after transplant, and 30 min at a time thereafter. Plants were given supplemental lighting up to to 14 hr day length and were not fertilized throughout the experiments.

**Data collection.** For each experiment, we measured plant height, leaf number, and chlorophyll content at four weeks after planting. Chlorophyll was measured using a SPAD 502 Plus (Spectrum Technologies, Inc. Aurora, IL); we recorded the mean of three measurements on the most recently emerged leaf. At the time of harvest, we counted total root nodules and collected 10 nodules from each plant to estimate per-nodule fresh weight. Shoots and roots were dried and weighed to determine the dry biomass of each plant.

**Rhizobium genomic data.** Full details are available in Riley et al. (6). Briefly, strains were grown as described above. Rhizobium genomic DNA was extracted and sequenced, followed by quality control and variant calling as described in Riley et al. (6). Variant data were hard-filtered using vcftools (v0.1.17, 8) to include variants with quality scores above 20. Multi-nucleotide polymorphisms were retained as single variants and sites that lacked genotype calls for more than 20% of the strains were removed. We included in the analyses only variants that had a minor allele frequency (MAF) of $\leq$ 5%. We found 491,227 variants represented in our genomes. After filtering for quality and frequency, 36,526 variants remained.

**Phenotypic analyses.** Within the R environment (9), we implemented linear mixed models (LMMs) using the R package lme4 (v1.1-27.1, 10) to test for G $\times$ E between 191 strains of rhizobia (G) and the two greenhouse experiments (E) for each plant line separately. We additionally partitioned G $\times$ E interactions into variance versus crossing effects using Cockerham's method (11, 12, 13). We square root-transformed all phenotypic variables to improve the normality of the data. For all phenotypic traits (shoot biomass, leaf number, plant height, chlorophyll content), the model included experiment and strain, and their https://doi.org/10.5281/zenodo.6712084 Zenodo: https://doi.org/10.5281/zenodo.6712084 Zenodo: https://doi.org/10.5281/zenodo.6712084 Zenodo: https://doi.org/10.5281/zenodo.6712084 Zenodo: https://doi.org/10.5281/zenodo.6



interaction as fixed effects. Rack was included as an additional random effect. From these models, we calculated estimate marginal means for each strain using the emmeans package (v1.4.1, 14).

Our experimental design, which featured two experiments nested within each host genotype, allows robust statistical estimation of G × E, but is underpowered to test for G × G interactions for partner quality phenotypes using ANOVA. Nevertheless a separate study (K. Heath, unpublished data) featuring a subset (N = 20) of the strains from the current study provides strong statistical support for G × G interactions for partner quality in this system (e.g., plant x strain G × G; $\chi^2$ = 83.3; p < 0.0001 for plant aboveground biomass; **Supp. Fig. S12**), similar to several other studies in this system (e.g., 15, 16, 17, 18, 19). We additionally calculated the broad-sense heritability ($H^2 = \frac{V_G}{V_P}$) for rhizobium strains within each experiment using LMMs in which response variables were square root-transformed, and both strain and rack were included as random effects. The significance of $V_G$ was assessed by comparing the full model to one in which the strain term was excluded, and a log-likelihood ratio test was performed between the two models to assess whether the proportion of variation explained (PVE) by the strain term was significant. Given the complex multivariate nature of our data at both the phenotypic and genomic levels, we focus the main text on shoot biomass, our core metric of partner quality; results for all other traits (i.e., leaf chlorophyll A, plant height, leaf number) can be found in the Supplementary Materials.

**Genome-wide association studies.** We performed multiple genome-wide association studies (GWAS) to identify rhizobium genomic variants associated with symbiotic partner quality. We conducted association tests for partner quality traits measured on plant hosts using a LMM approach to GWAS as implemented in the program GEMMA (v0.98.1, 20). Mapping analyses were performed separately for each of the four experiments, on standardized emmeans that corrected for the effects of rack (see above). GWAS relies on variants that are statistically distinguishable from one another (i.e., are not closely linked), and so we first identified genetic variants in strong linkage disequilibrium by splitting variants from all three rhizobia genomic elements into linkage groups based on the LD threshold $r_2 \geq 0.95$ using a customized script available at the Dryad repository associated with Epstein et al. (21), and picked a representative variant from each linkage group with the highest minor allele frequency and least missing data when ties were present (21). We ended up with 7,965 unlinked variants, 703 of which were on the chromosome, 2,955 were on pSymA, and 4,307 were on pSymB. We used selected variants to compute standardized kinship (k) matrices in GEMMA using the - gk 2 option, one for each genomic element, and performed the GWAS mapping using the lmm -4 option. In order to ensure our methods of calculating the k-matrix did not influence the results, we also conducted additional association tests for shoot biomass using k-matrices constructed for: 1) the entire genome (all three replicons together), and 2) for all variants, rather than those filtered by LD. However, our results were comparable regardless of the method used for calculating the k-matrix (**Supp. Fig. S13**), and thus, we only present results based on the initial method.

We assigned significance to particular variants using a permutation method which involves creating 1000 randomized datasets (i.e., strain identifiers randomized with respect to phenotypic means) and running the exact same LMM used for the non-randomized data on each in GEMMA. Therefore, all models include the same set of variants and the kinship matrix, the only difference between randomized and non-randomized models is the phenotype input file (.fam). After running our permutations, we tagged variants from the non-randomized run that fell above the 95% false discovery rate cut off based on the combined outputs of the randomized runs as being significant (21, 22). We summarized these significant variants to the gene-level by identifying the genes closest to or encompassing the variant using the intersect option in bedtools (v2.29.2, 23), excluding any intergenic regions. We additionally present the likelihood ratio test p-values generated by the standard GEMMA output files (.assoc) for both univariate LMMs (only one trait analyzed at a time) and multivariate LMMs (same trait analyzed across all four experiments). We calculated adjusted p-values for multiple tests using two approaches that control the strong-sense family-wise error rate: conventional Bonferroni correction (0.05 / 7,965 = 6.28 x $10^{-6}$) and the harmonic mean p-value (HMP) using the harmonicmeanp R package (v3.0, 24).

Finally, we were interested in the genetic basis of environmental-dependency (i.e., "G × E" genes), where the allelic effects varied among experiments within a particular host genotype due to either conditional neutrality (i.e., significant effects on the phenotype in one environment but not the other) or antagonistic allelic effects (i.e., significant effects in both environments, but in the opposite direction). Because mapping experiments suffer from false negatives (25), the lack of a significant association in one experiment does not rigorously identify patterns of conditional neutrality at the individual locus or at the global (whole genome) level (25, 26, 27). Thus we took multiple approaches to inspecting G × E in our association analyses. First, we used cross-experiment correlations between the estimated effects from each experiment (computed independently, see above) to visually assess the degree to which allelic effects were consistent across experiments (within a host genotype) and identify antagonistic allelic effects (loci with significant, but opposite, effects in the two experiments). Next, to test the global null hypothesis that the direction and magnitude of the estimated rhizobium allelic effects were consistent across the two experiments for a particular host (i.e., that allelic effects estimated in the two environments fell along the 1:1 line), we used a permutation test wherein we first resampled the estimated effects from the first experiment (i.e., DZA experiment I) with experimental error (standard deviation) and calculated the slope of their correlation, repeated this for 1000 permutations, then computed the probability of our observed slope given this simulated null distribution. Beyond this global test, to rigorously identify individual loci that contribute to the environmental response, we calculated plasticity for each strain (28, 29); plasticity was calculated as the natural log of the response ratio of the phenotype across experiments (e.g., $log(\frac{\text{shoot biomass exp. 3 (DZA)}}{\text{shoot bioamss exp. 1 (DZA)}})$; Lau et al. 30, Heath et al. 31) and mapped this trait separately for each of the two host genotypes.

**Candidate gene functional analyses.** To explore the biological interpretation of our various gene lists (e.g., A17-only, Experiment 1, Universal; see results), we used DAVID Bioinformatics Resources (v.6.8, 32) to test for significantly overrepresented UNIPROT keywords, GO terms, and KEGG pathways, as described in Sherman et al. (33). We report the results of enrichment analyses as “marginal” when the pre-FDR p-value associated with the EASE modified Fisher exact test was < 0.05 and “significant” when FDR-corrected p-value was < 0.05. We explored key pathways and genes implicated in particular gene sets using BioCyc (34, 35).

#### SUPP. DATASETS

- Dataset S1: **"SNPs_ann_ps_shoot.wREADME.xlsx"**
  Summary table for all variants significantly associated with shoot biomass in any of the four experiments. The first tab "SNPs_ann_ps_shoot" provides variant-level information, while the "README" tab provides a brief description of each column in the first tab.
- Dataset S2: **"genes_shoot_uniprot.wREADME.xlsx"**
  Summary table for all genes containing variants significantly associated with shoot biomass in any of the four experiments. The first tab "genes_shoot_uniprot" provides gene-level information, while the "README" tab provides a brief description of each column in the first tab.
- Dataset S3: **"SNPs_ann_ps_all.wREADME.xlsx"**
  Summary table for all variants significantly associated with one or more partner quality traits in any of the four experiments. The first tab "SNPs_ann_ps_all" provides variant-level information, while the "README" tab provides a brief description of each column in the first tab.
- Dataset S4: **"SNPs_ann_gene_all.wREADME.xlsx"**
  Summary table for all genes containing variants significantly associated with one or more partner quality traits. The first tab "SNPs_ann_gene_all" provides gene-level information, while the "README" tab provides a brief description of each column in the first tab.
- Dataset S5: **"DAVID_outputs_combined.wREADME.xlsx"**
  Summary table for all genes run through DAVID for which terms (GO, UNIPROT, KEGG pathways, et c.) were significantly (or marginally) enriched. The first tab "DAVID_outputs_combined" provides gene-level information, while the "README" tab provides a brief description of each column in the first tab.
- Dataset S6: **"plast_overlap_shoot.wREADME.xlsx"**
  Summary table for all genes containing variants significantly associated with either shoot biomass or plasticity for shoot biomass or both (i.e., overlapping). The first tab "plast_overlap_shoot" provides gene-level information, while the "README" tab provides a brief description of each column in the first tab.
- Dataset S7: **"genes_shoot.plast.wREADME.xlsx"**
  Summary table for all genes containing variants significantly associated with plasticity based on shoot biomass. The first tab "genes_shoot.plast" provides gene-level information, while the "README" tab provides a brief description of each column in the first tab.

Zenodo: https://doi.org/10.5281/zenodo.6712084

Zenodo: https://doi.org/10.5281/zenodo.6712084



Zenodo: https://doi.org/10.5281/zenodo.6712084



#### SUPP. FIGURES

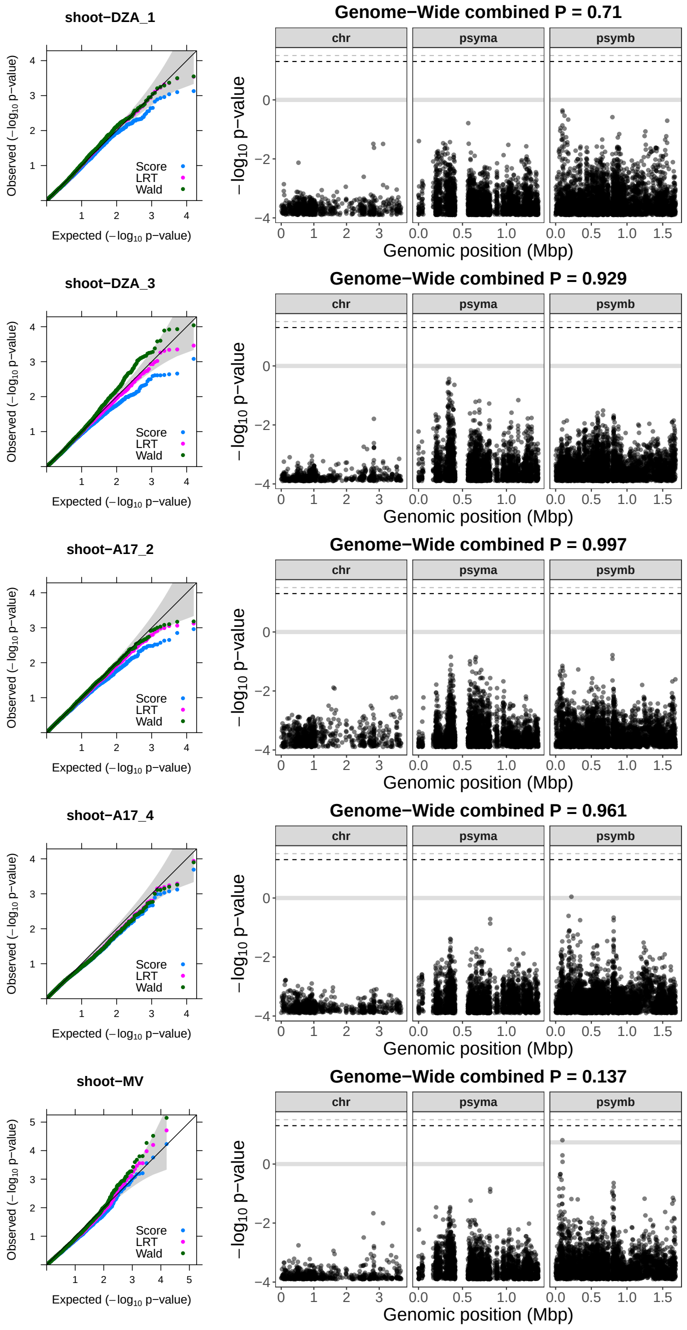


**Supp. Figure S1 Symbiotic extended phenotypes are governed by many variants with small effects.** QQ- (left) and Manhattan (right) plots of $\log_{10}$ p-values (based on likelihood ratio tests for variant associations with shoot biomass) calculated in GEMMA for each genomic region (chr = chromosome, psyma = symbiosis plasmid A, psymb = symbiosis plasmid B). The first four rows represent univariate linear mixed models (LMMs) completed for each experiment separately (DZA_1 = experiment 1 for DZA, DZA_3 = experiment 3 for DZA, et c.), while the bottom row represents the multivariate LMM for shoot biomass across the four experiments. Black dashed line represents the significance threshold based on Bonferroni correction (0.05/7,965), while the grey dashed line represents the harmonic mean p-value. The solid grey line represents the combined harmonic p-value for each genomic region.

https://doi.org/10.5281/zenodo.6712084 Zenodo: https://doi.org/10.5281/zenodo.6712084 Zenodo: https://doi.org/10.5281/zenodo.6712084 Zenodo: https://doi.org/10.5281/zenodo.6



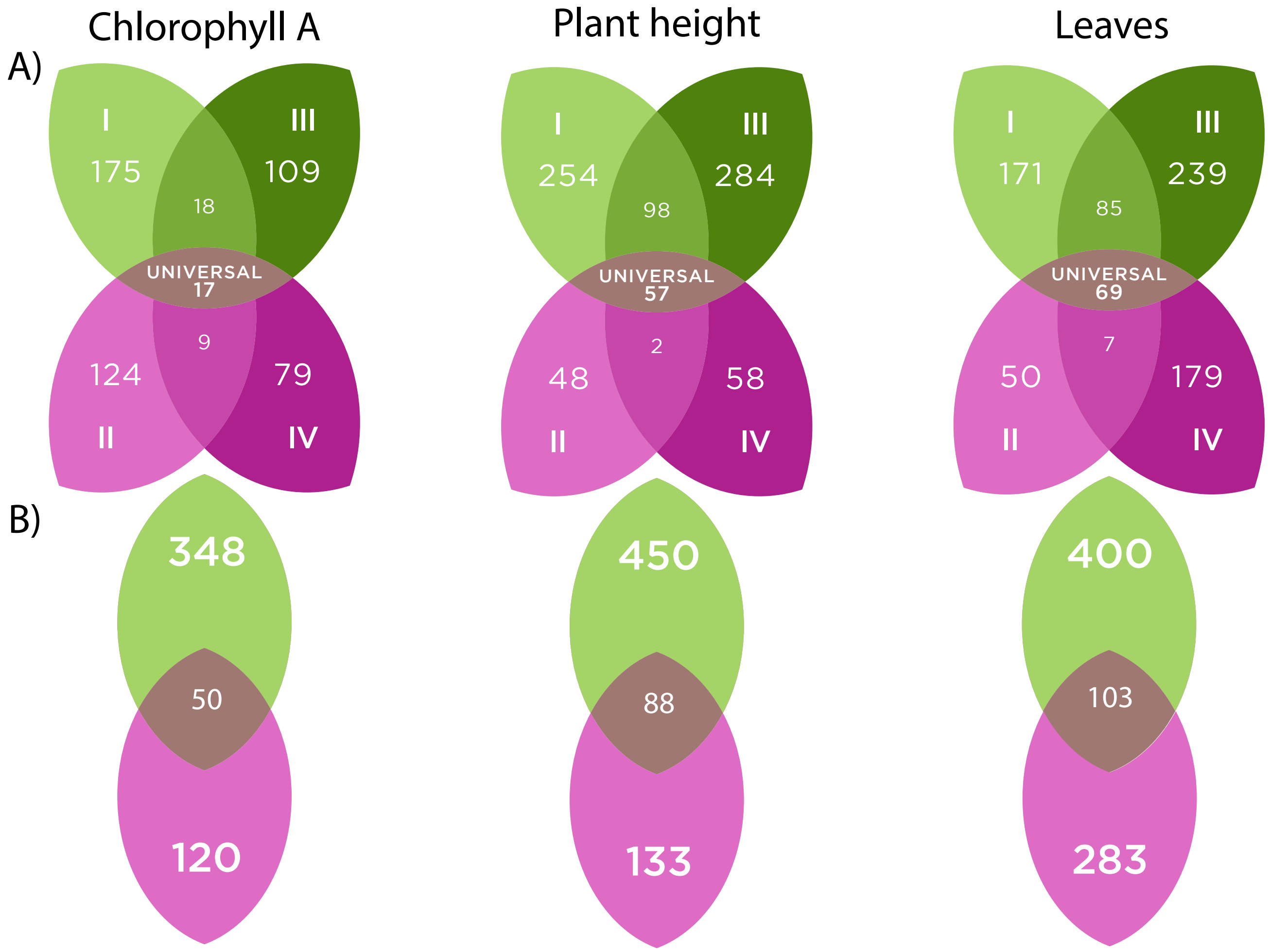


**Supp. Figure S2 Genes underlying G × E are prevalent across partner quality traits.** Venn diagrams showing number of rhizobium (*S. meliloti*) genes significantly associated with three other partner quality phenotypes for **A)** each of four separate mapping experiments or **B)** cross-experiment plasticity for either host genotype DZA in green (experiments I and III, in green) or A17 (experiments II and IV in pink). The mauve oval in the center represents either **A)** universal genes that contribute to trait variation in at least three of the four experiments or **B)** genes associated with cross-experiment trait plasticity in both host genotypes.

Zenodo: https://doi.org/10.5281/zenodo.6712084



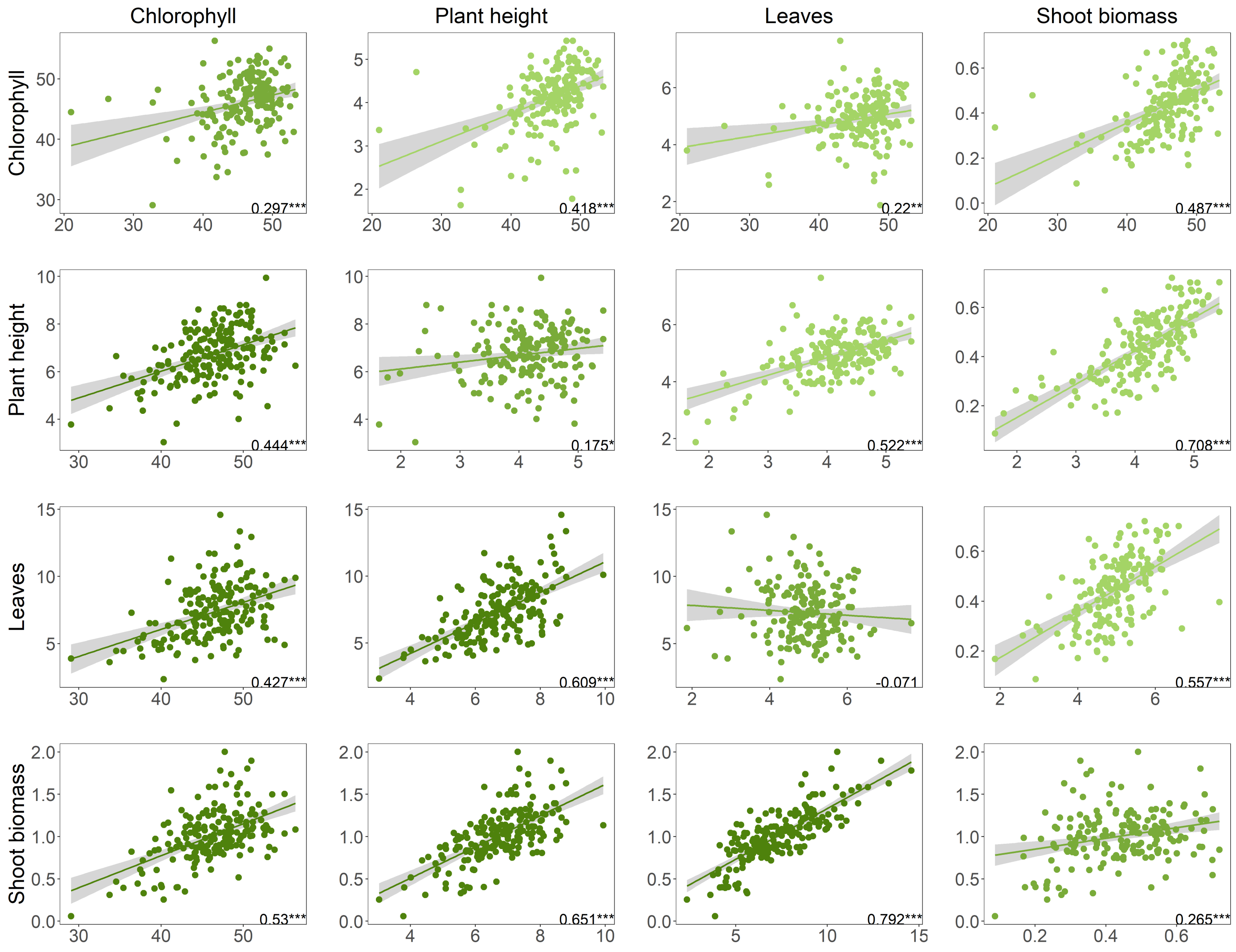


**Supp. Figure S3 Genetic correlations for traits measured on DZA.** Genetic correlations among traits within experiments (above or below diagonal) or between the same trait across experiments (along diagonal). Correlations based on estimated marginal means of each rhizobia strain corrected for rack on plant line DZA. Numbers in bottom right corners of each plot indicate Pearson correlation coefficients. Plots above and below the diagonal are for traits measured in experiments 1 and 3, respectively. Significance: $p < 0.001$ = '***'; $p < 0.01$ = '**'; $p < 0.05$ = '*'.

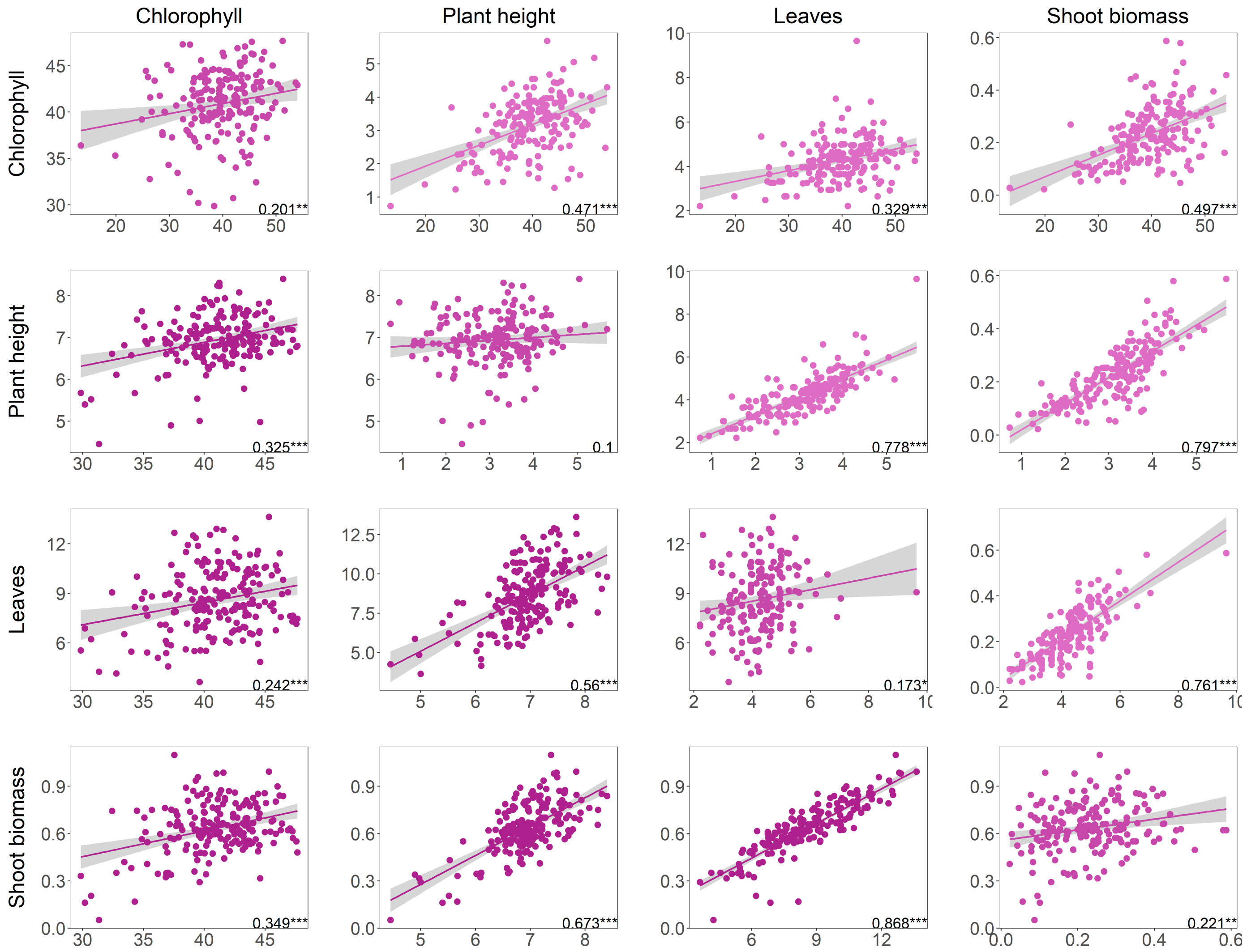


**Supp. Figure S4 Genetic correlations for traits measured on A17.** Genetic correlations among traits within experiments (above or below diagonal) or between the same trait across experiments (along diagonal). Correlations based on estimated marginal means of each rhizobia strain corrected for rack on plant line A17. Numbers in bottom right corners of each plot indicate Pearson correlation coefficients. Plots above and below the diagonal are for traits measured in experiments 2 and 4, respectively. Significance: p < 0.001 = '***'; p < 0.01 = '**'; p < 0.05 = '*'.

Zenodo: https://doi.org/10.5281/zenodo.6712084 Zenodo: https://doi.org/10.5281/zenodo.6712084 Zenodo: https://doi.org/10.5281/zenodo.6712084 https://doi.org/10.5281/zenodo.6712084

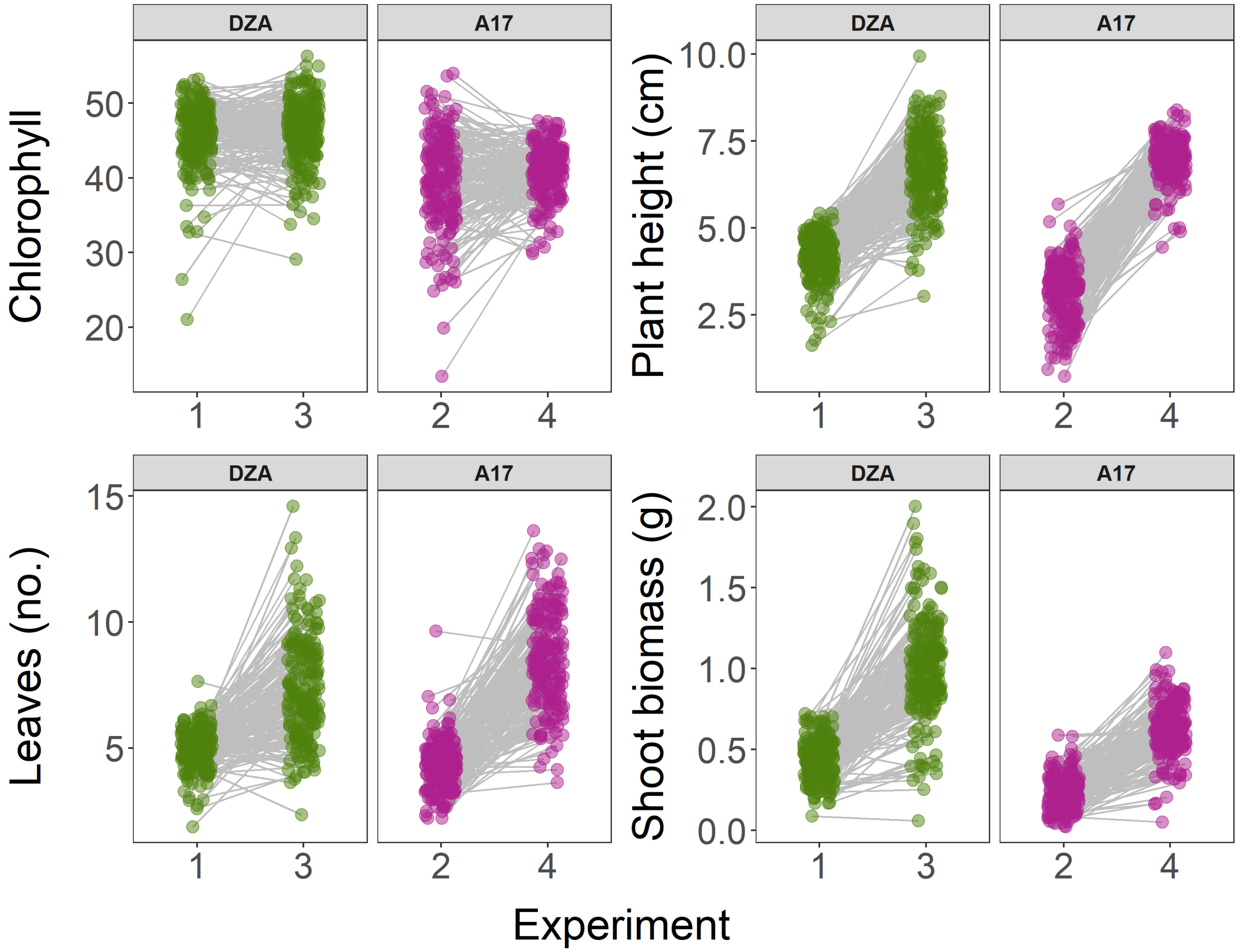


**Supp. Figure S5 Extensive G $\times$ E between experiments.** Reaction norms for partner quality traits across experiments. Data points represent estimated marginal means corrected for rack within each experiment. Points in green for means estimated on DZA, purple for A17.

Zenodo: https://doi.org/10.5281/zenodo.6712084 Zenodo: https://doi.org/10.5281/zenodo.6712084 Zenodo: https://doi.org/10.5281/zenodo.6712084 Zenodo: https://doi.org/10.5281/zenodo.6

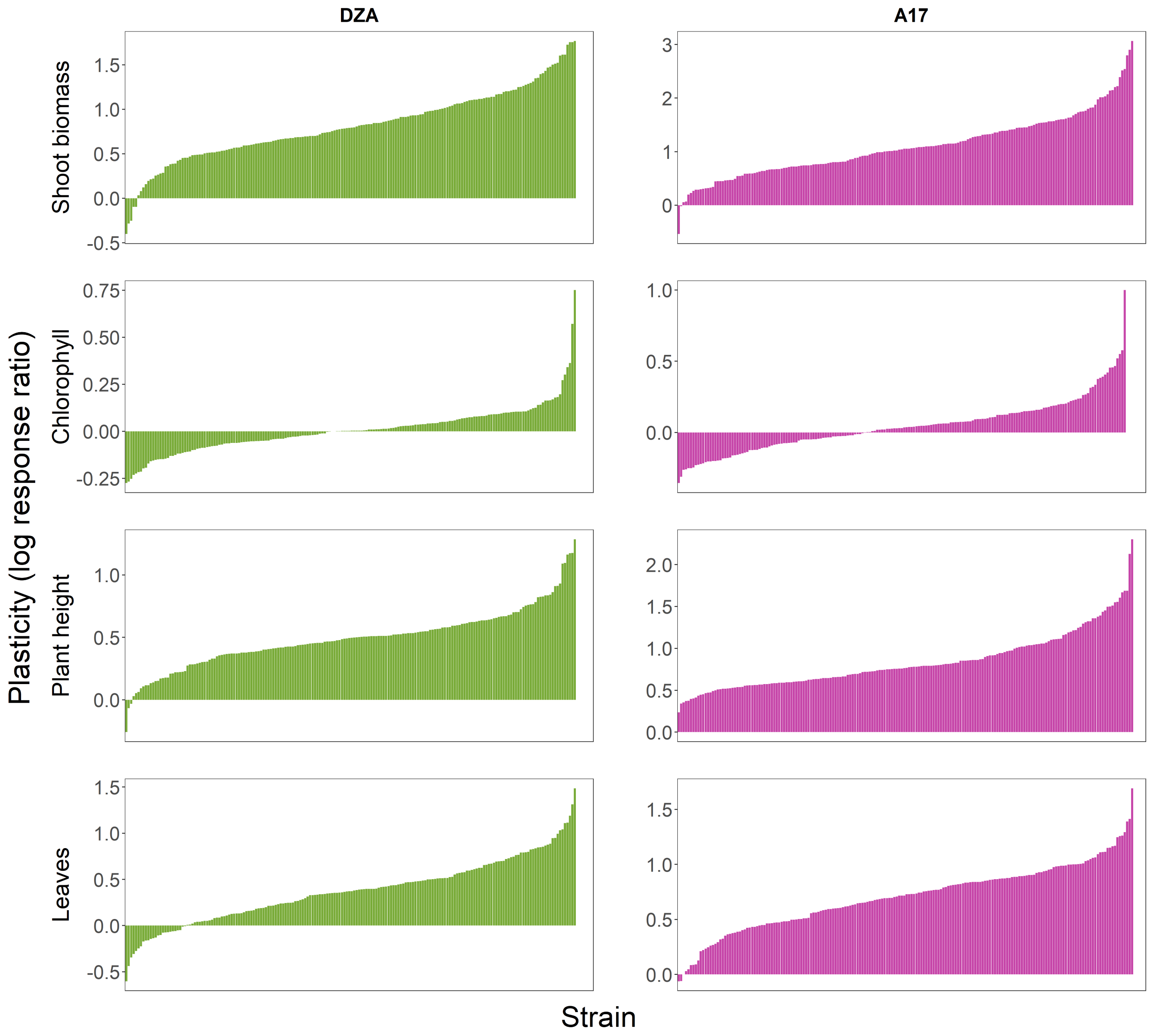


**Supp. Figure S6 Strains significantly varied in their response to different experiments.** Variation among *S. meliloti* strains in plasticity, calculated as the log response ratio for each trait between the two experiments with each host genotype (experiments 1 & 3 with DZA in green; experiments 2 & 4 with A17 in pink).

Zenodo: https://doi.org/10.5281/zenodo.6712084

Zenodo: https://doi.org/10.5281/zenodo.6712084 Zenodo: https://doi.org/10.5281/zenodo.6712084 Zenodo: https://doi.org/10.5281/zenodo.6712084 Zenodo: https://doi.org/10.5281/zenodo.6712084



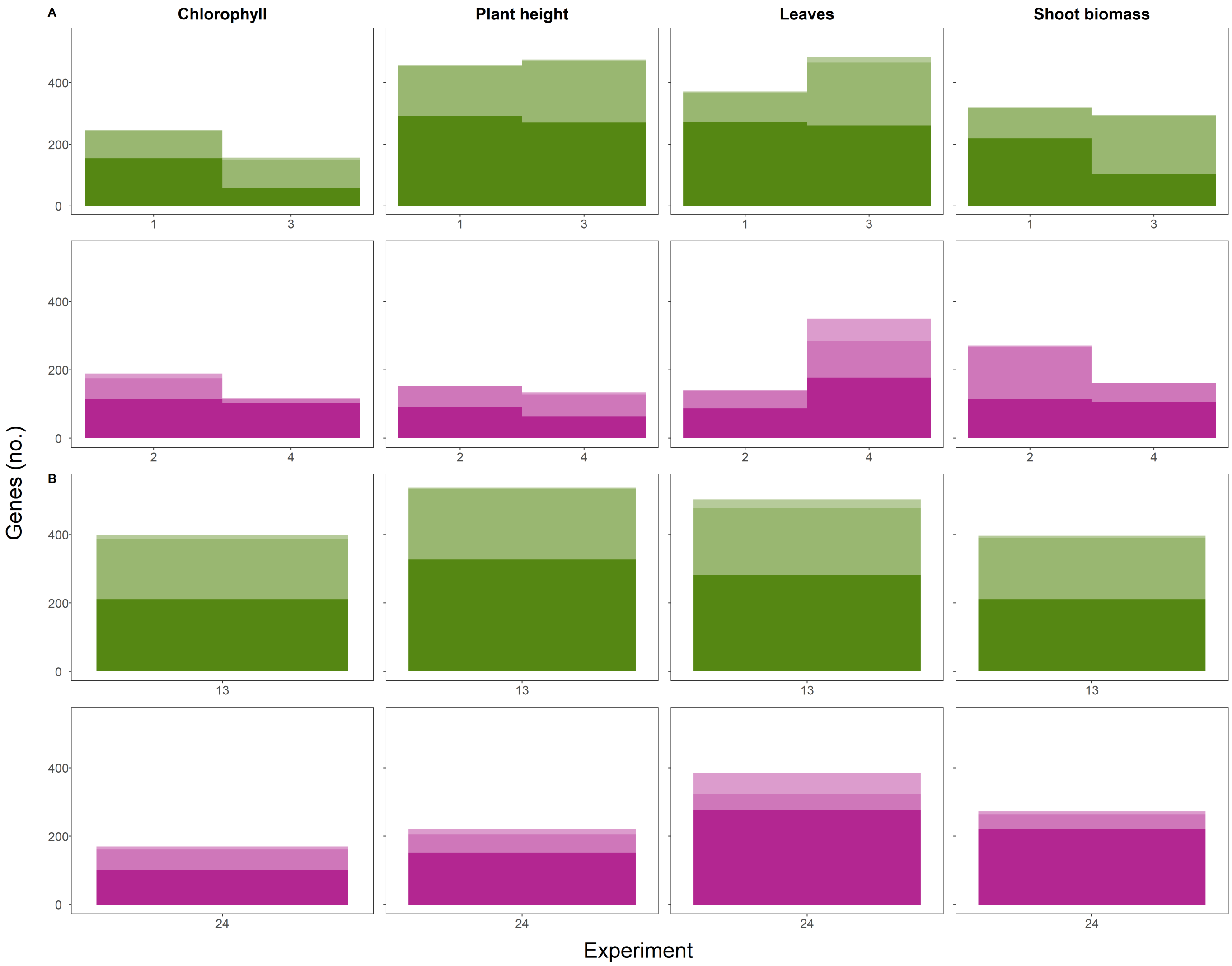


**Supp. Figure S7 Small-scale shifts of loci across genomic regions that contribute to partner quality variation.** Distribution of genomic locations (chromosome in lightest shade, pSymA in medium shade, and pSymB in darkest shade) for the *S. meliloti* loci significantly associated with partner quality phenotypes in each of four mapping experiments with either host genotype DZA (green) or host genotype A17 (pink). The plasticity panels **(B)** represent the genomic locations of rhizobium loci associated with the response of each trait across the two experiments for each host line (1-3 for DZA and 2-4 for A17).

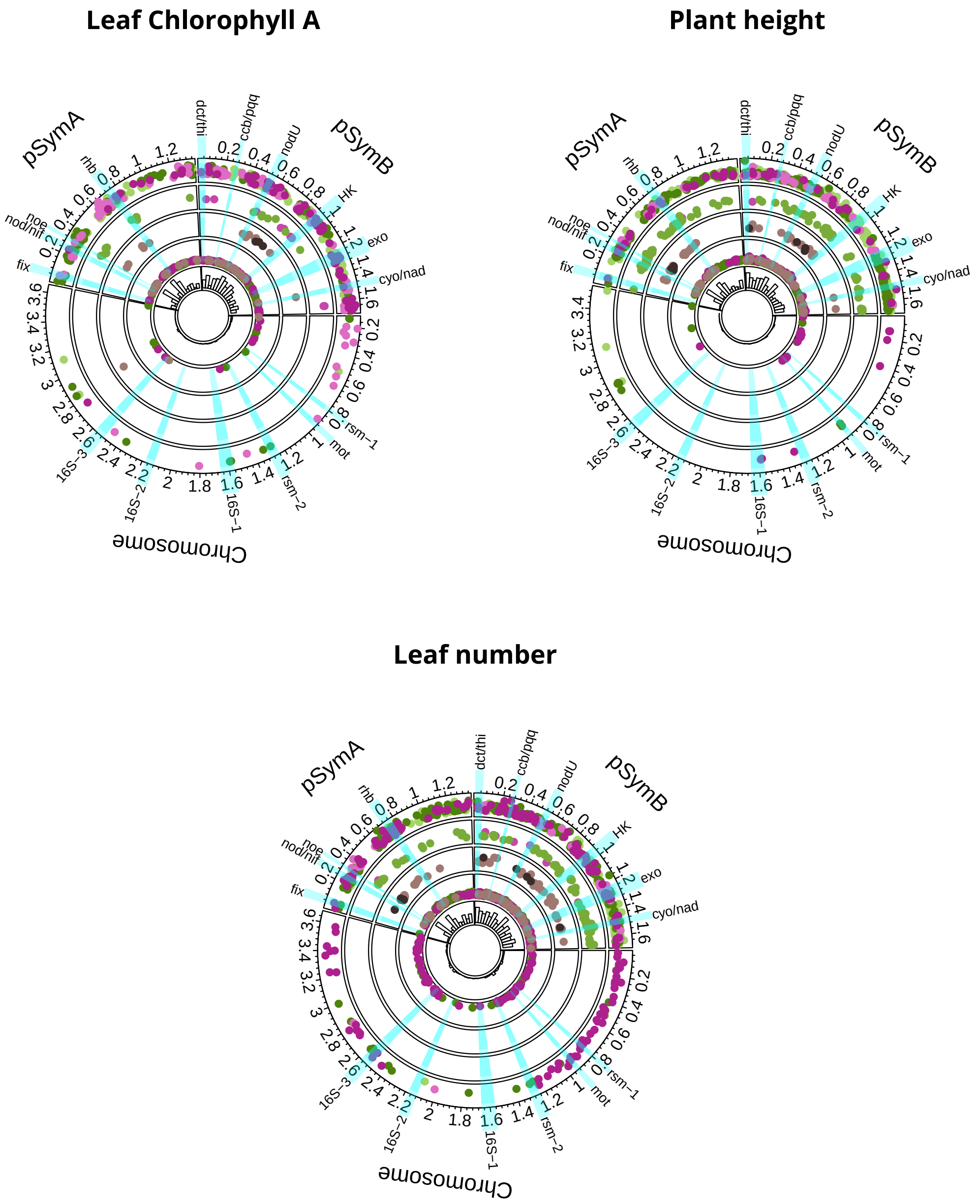


**Supp. Figure S8 Loci associated with partner quality are mostly limited to the symbiosis plasmids.** Circos plots showing positions of genes (dots) significantly associated with three additional partner quality traits. Each ring represents a different gene category, outermost to innermost: 1) G x E, 2) G x G, 3) partially universal and universal, 4) plasticity, while 5) depicts a histogram based on the total number of significant genes across 100 kbp-sized windows. The x- and y-axes for rings 1-4 represent genomic position (Mbp) and average absolute effect sizes of variants within each gene, respectively. The colours reflect categories in the Venn Diagrams: for rings 1, 2, and 4, genes associated with DZA-only traits are represented by shades of green, on A17-only with shades of purple, and both hosts in mauve (ring 4). For ring 3, genes associated with both hosts in more than three environments are represented in mauve (i.e., "partially universal"), and universal genes in black. Relevant loci are highlighted in blue, with abbreviations for clusters on the outer circle as follows: **rsm-1**: rsmD,E; ribF; groL; hisG,Z. **mot**: fliF,I,N,P,Q,R; flgB,D,F,G; motA,B; flhB. **rsm-2**: sppA; lptB; rpoN; raiA; ptsN; hrcA; rph; rdgB; ubiB; coaBC; iolB-E; cysK; rmsI; pyrF; queG, corA. **16S-1**: metB; rrf (5S rRNA); 16S rRNA; hrpB; hisA,H; addA,B; trxA; trpB; hpcH; gyrB; rho. **16S-2**: 16S rRNA; rrf (5S rRNA); oppD; glnQ; hppD; hmgA; maiA; purU; lpdA; modC; cobG,H,M. **16S-3**: grxC; ptsP; prmC; clpB; 16S rRNA; rrf (5S rRNA); tkt; deoC. **fix**: nnrU; norD,E; hemN; nirK; napA,E,F; fixG-L,P,Q,S; ccoN-Q; ric; nosR. **nod/nif**: nolF,G; nodA-C,D,D3,E,F,H,I,J,N; nifA,B,D,E,H,K,N,T,X; fabG; syrA; fdxB; fixA-C. **noe**: noeA,B; nodL; ccoN2,O2,P2,Q2; nodD2; groL. **rhb**: selB; fdhE; fdxH; fdnG; repA,B; katG; rhbC,D,F; basC; kdpB,F. **dct/thi**: urtA-C; dctD; thiC,O,S; mtnA; nspC; paaB,J,I,X. **ccb/pqq**: fghA; moxF; gfa; cbbX; rbcL; fba; tkt; pqqA,D,E. **nodU**: ugpC; ehuA-D; eutA,B; doeA,B; uxuA; galE; nodU. **HK** (housekeeping): alc; uraH; xdhA-C; guaD; pbpC; doeC; hutG,H; ltrA; phnC-E,N; gabD,T. **exo**: exsH; cueC-E; galE; exoA,F,H,I,K,L,M,O,P,Q,U,V,W,Y; thiD; nirD; nfeD, der; nnrU; bacA; map; glpK; xdhA; guaD; lldD. **cyo/nad**: cyoA-D; rfaL; minC,D; nadE; asnB.

https://doi.org/10.5281/zenodo.6712084 Zenodo: https://doi.org/10.5281/zenodo.6712084 Zenodo: https://doi.org/10.5281/zenodo.6712084 Zenodo: https://doi.org/10.5281/zenodo.6

Zenodo: https://doi.org/10.5281/zenodo.6712084 Zenodo: https://doi.org/10.5281/zenodo.6712084 Zenodo: https://doi.org/10.5281/zenodo.6712084



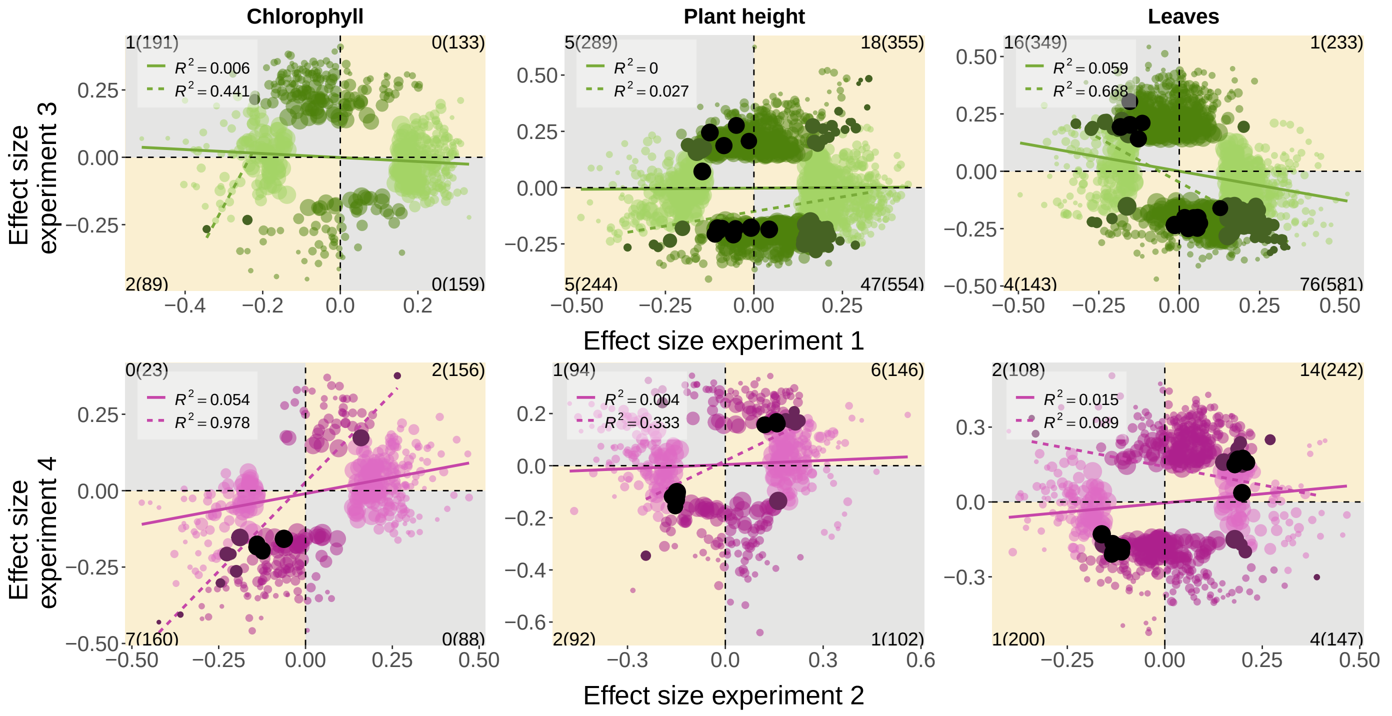


**Supp. Figure S9 Extensive G × E revealed at the variant-level.** Variant-level G × E for partner quality loci. Shown are correlations between the estimated effects of individual *S. meliloti* loci on three difference partner quality metrics (from GWAS) in each of two experiments for either host DZA (green) and A17 (pink). Only allelic effects that were significant in one (lighter colours) or both (dark points) environments are shown, while black dots represent nearly universal variants, i.e., associated with the same trait in three experiments. Linear relationships and $R^2$ values are depicted for all significant variants (solid coloured line) or variants significant in both experiments (dotted coloured line). Variant counts for each quadrant are shown in the corners of each plot (variants significant in both experiments, followed by all variants in parentheses).

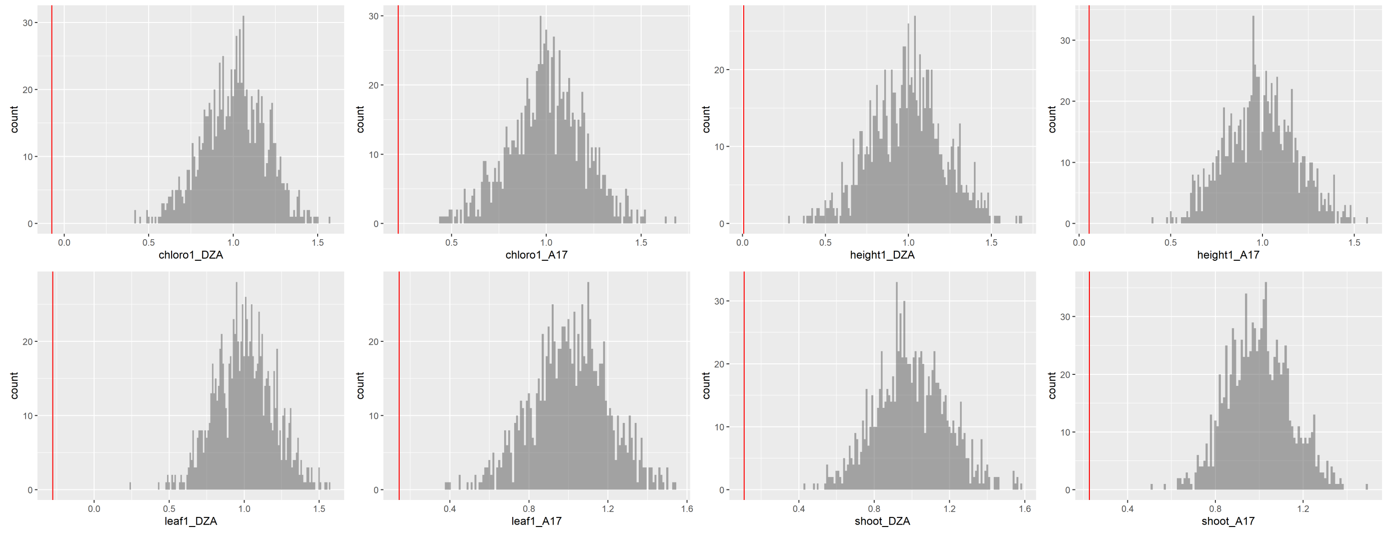


**Supp. Figure S10 Global analyses show that allelic effects are significantly different across experiments.** Distributions of slopes calculated by resampling the estimated effects from one experiment (e.g., DZA experiment I) with experimental error (standard deviation), regressing against the observed effects in that experiment, and repeating 1000 times. Red lines depict observed slopes when estimated effects were regressed between experiments.

https://doi.org/10.5281/zenodo.6712084 Zenodo: https://doi.org/10.5281/zenodo.6712084 Zenodo: https://doi.org/10.5281/zenodo.6712084 Zenodo: https://doi.org/10.5281/zenodo.6712084 Zenodo: https://doi.org/10.5281/zenodo.6



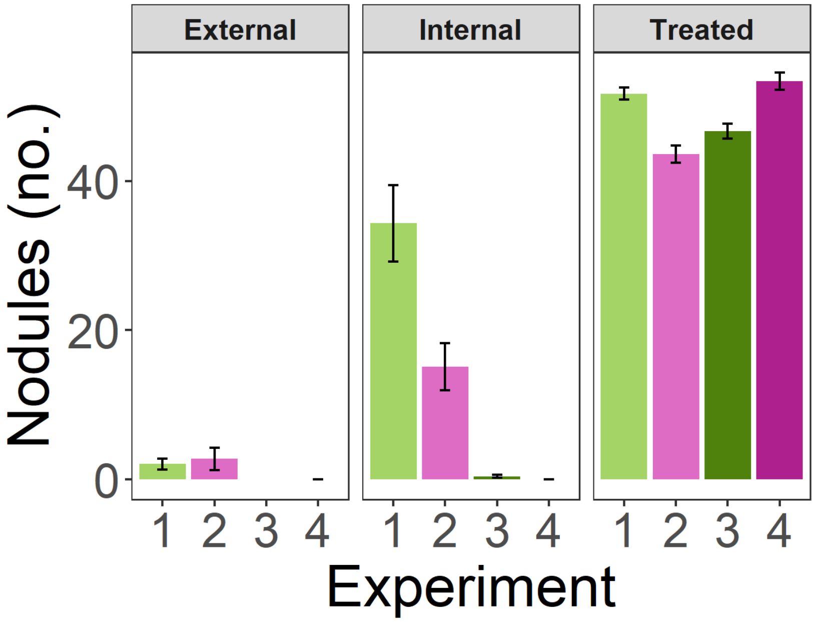


**Supp. Figure S11 Contamination was minimal across experiments.** Nodule number for inoculated (i.e., Treated) versus Internal and External uninoculated control plants in four experiments: Experiments 1 and 3 used host genotype DZA (shades of green) while 2 and 4 used genotype A17 (shades of pink).

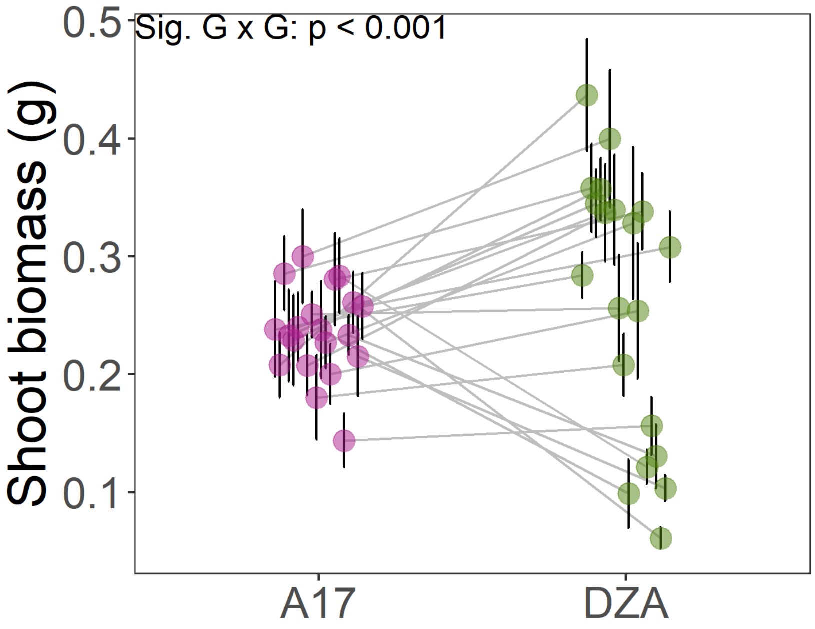


**Supp. Figure S12 Significant Genotype-by-genotype (G × G) interactions between two host genotypes (A17 and DZA) and 20 *S. meliloti* strains.** All 20 strains were included in the 191 strains used in current paper. Both hosts were grown together in a single experiment (Heath et al., unpublished data). Type III ANOVA based on a linear mixed model that corrected for rack and researcher was used to test for the main effects of rhizobium strain ($\chi^2$ = 94.891, p < 0.001), host genotype ($\chi^2$ = 0.553, p = 0.457), and strain-by-host (G x G) interaction ($\chi^2$ = 83.272, p < 0.001) on plant shoot biomass.

Zenodo: https://doi.org/10.5281/zenodo.6712084 Zenodo: https://doi.org/10.5281/zenodo.6712084 Zenodo: https://doi.org/10.5281/zenodo.6712084 Zenodo: https://doi.org/10.5281/zenodo.6



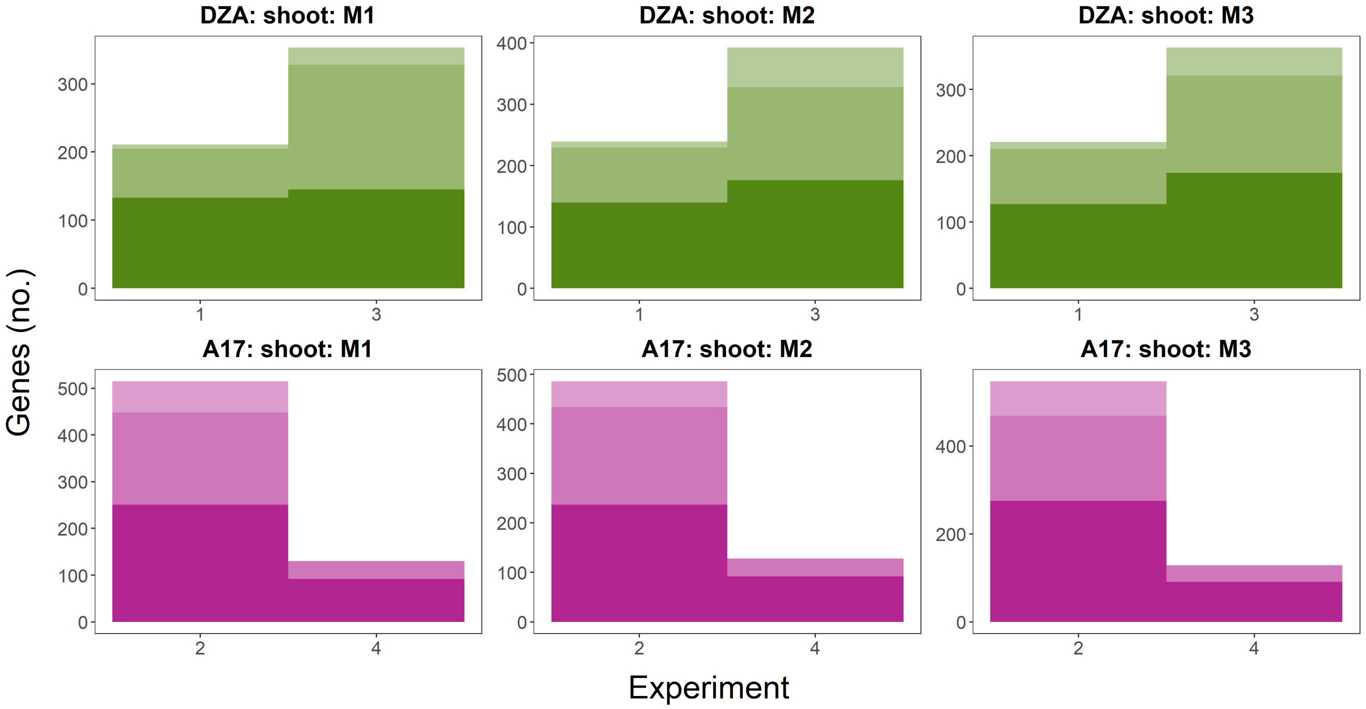


**Supp. Figure S13 Minimal differences in the number of significant genes across three different methods.** To ensure the significant variants identified in our study did not depend on the computational methods used (i.e., how the k-matrix was computed), we compared the number of genes tagged by variants significantly associated with shoot biomass in all four experiments and both plant lines (top row in shades of green = DZA; bottom row in shades of pink = A17) for three separate methods: **M1)** k-matrix calculated for each genomic region separately, only unlinked variants included as input; **M2)** k-matrix calculated for the whole genome, only unlinked variants included as input; and **M3)** k-matrix calculated for the whole genome, linked variants included as input. The different shades represent whether genes were located on the chromosome (lightest), pSymA (medium), and pSymB (darkest).
